# Disrupted Nitric Oxide Homeostasis Impacts Fertility Through Multiple Processes Including Protein Quality Control

**DOI:** 10.1101/2024.07.30.605885

**Authors:** Patrick Treffon, Elizabeth Vierling

**Affiliations:** Department of Biochemistry and Molecular Biology, University of Massachusetts Amherst, Amherst, MA, USA

**Keywords:** nitric oxide_1_, S-nitrosoglutathione reductase (GSNOR)_2_, fertility_3_, pistils_4_, S-nitrosoproteome_5_, UDP-glycosyltransferase_6_, Argonaute proteins_7_, 26S proteasome_8_

## Abstract

Plant fertility is fundamental to plant survival and requires the coordinated interaction of developmental pathways and signaling molecules. Nitric oxide (NO) is a small gaseous signaling molecule that plays crucial roles in plant fertility as well as other developmental processes and stress responses. NO influences biological processes through S-nitrosation, the posttranslational modification of protein cysteines to S-nitrosocysteine (R-SNO). NO homeostasis is controlled by S-nitrosoglutathione reductase (GSNOR), which reduces S-nitrosoglutathione (GSNO), the major form of NO in cells. GSNOR mutants (*hot5-2/gsnor1*) have defects in female gametophyte development along with elevated levels of reactive nitrogen species and R-SNOs. To better understand the fertility defects in *hot5-2* we investigated the *in vivo* nitrosoproteome of floral tissues coupled with quantitative proteomics of pistils. To identify protein-SNOs we employed, for the first time in plants, an organomercury-based method that involves direct reaction with S-nitrosocysteine, enabling specific identification of S-nitrosocysteine–containing peptides and S-nitrosated proteins. We identified 1102 endogenously S-nitrosated proteins in floral tissues, of which 1049 were unique to *hot5-2.* Among the identified proteins, 728 were novel S-nitrosation targets. Notably, specific UGT-glycosyltransferases and argonaute proteins are S-nitrosated in floral tissues and differentially regulated in pistils. We also discovered S-nitrosation of proteins of the 26S proteasome together with increased abundance of proteasomal components and enhanced trypsin-like proteasomal activity in *hot5*-*2* pistils. Our data establish a new method for nitrosoprotein detection in plants, expand knowledge of the plant S-nitrosoproteome, and suggest that nitro-oxidative modification and NO homeostasis are critical to protein quality control in reproductive tissues.

## 1 Introduction

Nitric Oxide (NO), a gaseous signaling molecule, has emerged as a key player in mediating diverse physiological responses in plants (Lee et al., 2008; Kolbert et al., 2021; Liao et al., 2023). NO belongs to the group of reactive nitrogen species (RNS), molecules that contain nitrogen and typically have one or more unpaired electrons, making them highly reactive and capable of participating in multiple chemical reactions (Mock and Dietz, 2016; Treffon and Vierling, 2022). NO also reacts with the tripeptide glutathione (GSH) to form S-nitrosoglutathione (GSNO), which acts as a mobile NO reservoir in plants (Zaffagnini et al., 2013). In general, RNS serve in biological systems as signaling molecules and influence multiple cellular processes. However, an imbalance or excessive production of RNS can lead to nitro-oxidative stress that can result in cellular damage and ultimately impact plant health negatively (Terron-Camero et al., 2020; Wani et al., 2021). The signaling effects of RNS are primarily the result of post-translational modifications of protein Cys residues via S-nitrosation (R-SNO), the covalent attachment of a NO-group to cysteine residues of proteins. S-nitrosation can impact protein function, localization and interactions (Guerra et al., 2016; Jedelská et al., 2020; Corpas et al., 2022).

A major component regulating GSNO and therefore RNS homeostasis is the NADH-dependent cytosolic enzyme S-nitrosoglutathione reductase (GSNOR), which modulates the effects of RNS through catabolism of GSNO (Li et al., 2020). It is a highly conserved enzyme across plant and animal kingdoms and GSNOR-null mutants in *A. thaliana* (*hot5*-*2/gsnor1-3*) exhibit a broad spectrum of phenotypic defects including an elevated level of NO species, heat sensitivity, compromised immune-response, increased reproductive shoots and profoundly reduced fertility (Lee et al., 2008; Xu et al., 2013; Chae et al., 2021; Wang et al., 2021; Wang et al., 2024). Plant reproduction is a complex process important for the continuation of species and crop production. It includes a series of tightly coordinated events, from pollen and female development to fertilization and seed formation. Female gametophyte (FG) development, a crucial stage in plant reproduction, involves cell divisions leading to formation of the egg cell, along with antipodal, synergid and central cells within the ovule (Drews and Koltunow, 2011). The absence of GSNOR in *hot5-2* mutants leads to the accumulation of RNS, particularly in the sporophytic nucellar tissue surrounding the developing female gametophyte. This disregulated NO homeostasis hinders the efflux of auxin into the developing FG, and the reduced auxin levels correlate with FG degeneration and decreased seed yield (Wang et al., 2021; Wang et al., 2024). *hot5-2* mutants also show delayed appearance or mispositioning of different FG cell types (Wang et al. 2024). However, how elevated RNS levels and increased R-SNOs contribute to these FG phenotypic defects in GSNOR mutants remains to be elucidated.

Determining what proteins are nitrosated in the absence of GSNOR is a first step toward defining molecular mechanisms by which disrupted NO homeostasis leads to FG defects. However, accessing the global S-nitrosoproteome is challenging because of the labile nature of the R-SNO bond as well as a relatively low abundance of endogenously S-nitrosated proteins (Astier et al., 2012; Lamotte et al., 2015). The biotin-switch assay (BSA) and its derivative assays (S-nitrosothiol-Resin Assisted Capture or SNO-RAC) combined with mass spectrometry (MS) are commonly used to identify S-nitrosated proteins in different species (Forrester et al., 2009; Forrester et al., 2009; Hu et al., 2015). Both the BSA and SNO-RAC methods depend on blocking free thiols by alkylation followed by ascorbate-mediated conversion of S-NO groups to free thiols. The newly reduced, free thiols are then chemically conjugated to an affinity tag (e.g. biotin-HPDP (*N*-[6-(Biotinamido)hexyl]-3′-(2′-pyridyldithio)propionamide); thiopropyl-sepharose) and purified, and the enriched S-nitrosated proteins analyzed by Western blotting or mass spectrometry. However, several reports question the specificity of the ascorbate-mediated reduction of the R-SNO bond that constitutes a critical step in these procedures and that may lead to false-positive identification of R-SNOs (Landino et al., 2006; Monteiro et al., 2007; Giustarini et al., 2008; Wang et al., 2008). For the first time in plants, we employed an organomercury-based approach to identify S-nitrosated proteins. This method involves direct reaction of an organomercury-affinity matrix (*p*-aminophenylmercuric-acetate coupled to agarose) with R-SNOs, as opposed to the indirect methods requiring prior SNO reduction by ascorbate. The organomercury method takes advantage of the high affinity of organomercury towards S-nitrosated cysteine residues, eliminating the problematic ascorbate reduction step. Starting with protein extracts (1) the method involves: (2) blocking reduced cysteines with methyl methanethiosulfonate (MMTS), (3) direct capture of S-nitrosated proteins or peptides with *p*-aminophenylmercuric-acetate coupled to agarose beads, release of peptides after tryptic digestion (4) with or without mild performic acid (5), and (6) liquid chromatography/tandem MS analysis (Doulias and Gould, 2018). An important control involves parallel samples in which the SNO is cleaved by UV photolysis prior to reaction with *p*-aminophenylmercuric-acetate. The performic acid-mediated release step also oxidizes cysteine thiols to sulfonic acid, creating a unique MS signature that permits site-specific identification of the modified cysteines.

We profiled the S-nitrosoproteome of floral tissues of the *hot5-2* GSNOR null mutant and wild type (WT) because the endogenous increase in RNS species in the mutant (Lee et al., 2008; Xu et al., 2013) predicted higher levels of protein S-nitrosation compared to WT plants. A previous nitrosoproteome study of WT and *hot5-2* seedlings indeed uncovered 926 nitrosated proteins (Hu et al., 2015). By identifying alterations in the floral S-nitrosoproteome, we aimed to gain insights into the mechanisms behind the defects in *hot5-2* FG development. We coupled this analysis with total label-free quantitative (LFQ) proteomics of pistil proteins compared to sepal plus petal proteins (SePel) and leaf proteins to further define protein differences connected to the *hot5-2* fertility defect. We identified that proteins belonging to UGT-glycosyltransferases are highly upregulated in different *hot5-2* tissues, whereas argonaute (AGO) proteins are downregulated in *hot5-2* pistils. In addition, based on the S-nitrosoproteome combined with the quantitative protein abundance profiling of specific floral tissues, we found targeted S-nitrosation of proteins associated with the 26S proteasome together with elevated protein levels of proteasomal components and increased proteasome specific activity within *hot5-2* pistils. These data suggest a regulatory role for protein S-nitrosation in modulating protein quality control in plant reproductive tissues and provide valuable groundwork for future investigations into mechanisms impacted by nitro-oxidative modification.

## 2 Materials and Methods

### 2.1 Plant material and growth conditions

*Arabidopsis thaliana* Col-0 WT and *hot5-2* (GABI_315D11) plants were grown on soil at 60-100 µmol m^-2^ s^-1^ with a 16h light/8h dark schedule and 22 °C/18 °C.

### 2.2 Organomercury-resin-assisted capture of S-nitrosated proteins

The organomercury-resin-assisted capture (MRC) of S-nitrosoproteins was done with modifications as described (Doulias and Gould, 2018). Entire inflorescences pre-anthesis, corresponding to flower developmental (FD) stages of up to 12/13 (prefertilization) of soil grown WT and *hot5*-*2* plants were harvested, shock frozen in liquid N_2_ and ground to a fine powder. 5 mL of plant material was used to extract proteins using a 1:1 ratio of extraction buffer (250 mM Hepes pH 7.7, 1 mM diethylenetriaminepentaacetic dianhydride (DTPA), 0.1 mM neocuproine, 40mM *S*-Methyl methanethiosulfonate (MMTS), 1% TritonX-100). After 5 min of vortexing, samples were centrifuged at 16.000 x g at 4 °C for 15 min to separate cell debris from soluble protein. Centrifugation was repeated, protein concentration measured using the BCA assay (Thermo Scientific, USA), and 10 mg of total protein per sample was used for subsequent S-nitrosoprotein enrichment. For photolysis negative controls 0.1 M mannitol and 5 mM MMTS were added to 10 mg of protein homogenate, vortexed briefly and placed into borosilicate glass vials. Vials were placed on ice and illuminated for 7 min in “time mode” using a UV-crosslinker with a 254 nm light source (Stratalinker Model 1800, Stratagene, USA). Each sample had a corresponding photolysis negative control. Proteins from all samples were obtained after acetone precipitation and centrifugation at 4500 x g for 15 min at 4 °C by resuspending in blocking buffer (250 mM Hepes pH 7.7, 1 mM DTPA, 0.1 mM neocuproine, 50 mM MMTS, 2.5 % SDS). Samples were alkylated for 35 min in the dark in a 50 °C water bath with frequent vortexing, followed by another round of acetone precipitation to remove any non-reacted MMTS. Proteins were resuspended to 0.5 mg/ml in loading buffer (250 mM MES pH 6.0, 1 mM DTPA, 1 % SDS).

Organo-mercury resin (*p*-aminophenylmercuric-acetate coupled to agarose) was synthesized as described (Doulias and Gould, 2018). After alkylation, proteins were loaded onto 15 ml of activated organo-mercury resin slurry (50 %) and incubated for 60 min at RT in the dark to allow binding of S-nitrosated proteins to the resin. Following incubation, columns were washed with 50 column volumes (CV) of 50 mM Tris-HCl pH 7.5, 0.3 M NaCl, 0.5 % SDS; 50 CV of 50 mM Tris-HCl pH 7.5, 0.3 M NaCl, 0.05 % SDS; 50 CV of 50 mM Tris-HCl pH 7.5, 0.3 M NaCl, 1 % Triton X-100; 50 CV of 50 mM Tris-HCl pH 7.5, 0.3 M NaCl, 0.1 % Triton X-100, 0.1 M urea and 200 CV of H_2_O. For on-column trypsin digest, columns were pre-equilibrated with 5 CV of 0.1 M NH_4_CO_3_ and then loaded with 1 CV of 0.1 M NH_4_CO_3_ containing 1 µg/ml trypsin gold (Promega V5280, Promega, USA). The flow-through containing the digested peptides of S-nitrosated proteins (elution 1) was collected in glass tubes after incubation overnight at room temperature in the dark. The remaining S-nitrosated peptides that were bound to the organo-mercury resin were washed with 40 CV of 1 M NH_4_CO_3_, 0.3 M NaCl; 40 CV of 1 M NH_4_CO_3_; 40 CV of 0.1 M NH_4_CO_3_ and 200 CV of H_2_O. Bound peptides were oxidatively released (elution 2) by incubation with 1 CV of 1 % performic acid for 45 min at RT and collected in glass tubes. Following lyophilization of the samples and resuspension in 300 µl of 0.1 % formic acid, sample volume was reduced to 30 µl using a vacuum concentrator (Thermo Savant SPD111V, Thermo Scientific, USA) and centrifuged for 20 min at 16,000 x g at 4°C to remove any precipitate. Peptides were desalted using C18 tips as recommended by the manufacturer (Pierce C18 Tips, Thermo Scientific, USA) and resuspended in 15 µl of 1 % formic acid.

For the LC-MS\MS run, a 5 µL injection was loaded by a Thermo Easy nLC 1000 UPLC onto a 2 cm trapping column and desalted with 12 µL mobile phase A (0.1% formic acid in water). Peptides were eluted at 300 nL/min on to a 75 um x 15 cm RSLC column (Thermo) using a linear gradient of 5-35% mobile phase B (0.1% formic acid in acetonitrile) over 90 min. Ions were introduced by positive ESI using a stainless steel capillary at 2.1 kV into a Thermo Orbitrap Fusion tribrid mass spectrometer. Mass spectra were acquired over m/z 350-2000 at 120,000 resolution (m/z 200) and 1 sec cycle time with an automatic gain target (AGC) of 1e6, and data-dependent acquisition selected the top speed most abundant precursor ions for tandem mass spectrometry by HCD fragmentation using an isolation width of 2 Da and automatic gain settings. Peptides were fragmented with a normalized collision energy of 27 %, and fragment spectra acquired in the linear ion trap. Targeted mass exclusion was activated to avoid MS/MS of ions previously identified as Rubisco or keratin peptides.

For elution samples 1, which represent peptides from S-nitrosated proteins that were not bound to the column by S-nitrosocysteine peptides, mass spectra were searched against the Uniprot *A. thaliana* database (UP000006548_3702 and UP000006548_3702_additional, downloaded 07/2019) using MaxQuant version 1.6.17.0 with a 1% FDR at the peptide and protein level, peptides with a minimum length of seven amino acids with N-terminal acetylation and methionine oxidation as modifications. Enzyme specificity was set as C-terminal to arginine and lysine using trypsin as protease and a maximum of two missed cleavages were allowed in the database search. The maximum mass tolerance for precursor and fragment ions was 4.5 ppm and 20 ppm, respectively, with second peptides enabled. Label-free quantification (LFQ) was performed with the MaxLFQ algorithm using a minimum ratio count of 2. For elution samples 2 containing S-nitrosocysteine peptides, similar MaxQuant settings (version 1.6.17.0) as mentioned above were used with the exception that N-terminal acetylation, methionine oxidation and cysteine trioxidation were set as variable modifications and up to 5 modifications per peptide were allowed.

The identified protein groups generated by the MaxQuant program were uploaded to the Perseus program version 1.6.15.0 (Tyanova et al., 2016). Site only, reverse, and contaminant peptides were removed from the dataset and only protein groups identified in two of the three biological replicates (in at least one group) were further processed. Invalid values were excluded, and empty columns removed. Missing LFQ values were imputed using the default Perseus settings assuming a normal distribution. The Hawaii plot function with default parameters was used to identify S-nitrosated proteins that were significantly enriched in each sample compared to all other samples. For elution samples 2 containing the information for site-specific S-nitroso peptides, the MaxQuant output file for tri-oxidized cysteines (R-SO_3_H) was used. Gene set enrichment analysis was done using the online tool ShinyGO (Version 0.80, http://ge-lab.org/go/) with the KEGG pathway database selected (Ge et al., 2020). The FDR cutoff was set to 0.05 with minimum and maximum pathway sizes of 2 and 2000, respectively. MapMan analysis was conducted to provide a graphical overview of metabolic and regulatory pathways for the detected S-nitrosated proteins (Thimm et al., 2004; Usadel et al., 2009). For that, AGI identifiers for each protein were mapped against the TAIR10 mapping database.

### 2.3 LFQ proteomics of pistils

Pistils of *hot5*-*2* and WT at stages before pollination (FD 12-13; Fig. S4) were harvested and immediately shock frozen in liquid N_2_. Around 50 pistils per genotype were pooled as one biological replicate with a total of three biological replicates per genotype. Proteins were extracted using diluted HENS buffer [25mM HEPES pH 7.7, 1mM EDTA, 2.5 % SDS, protease inhibitor cocktail (Pierce A32955)], TCA precipitated and resuspended in the same buffer. After BCA protein determination, 50 µg of each whole cell lysate was run on a 4-20% SDS-PAGE for 15 min to separate proteins from lower molecular weight contaminants. The entire protein region was excised from the gel and subjected to in-gel trypsin digestion after reduction with dithiothreitol and alkylation with iodoacetamide as described in (Treffon et al., 2021). Peptides eluted from the gel were lyophilized and re-suspended in 50µL of 5% acetonitrile [0.1% (v/v) formic acid (FA)]. For each biological replicate, three technical replicates were measured by LC-MS/MS. A 2 µL injection was loaded by a Waters NanoAcquity UPLC in 5% acetonitrile (0.1% formic acid) at 4.0 µL/min for 4.0 min onto a 100 µm I.D. fused-silica pre-column packed with 2 cm of 5 µm (200Å) Magic C18AQ (Bruker-Michrom). Peptides were eluted at 300 nL/min from a 75 µm I.D. gravity-pulled analytical column packed with 25 cm of 3 µm (100Å) Magic C18AQ particles using a linear gradient from 5-35% of mobile phase B (acetonitrile + 0.1% formic acid) in mobile phase A (water + 0.1% formic acid) over 90 min. Ions were introduced by positive electrospray ionization via liquid junction at 1.4kV into a Thermo Scientific Q Exactive hybrid mass spectrometer. Mass spectra were acquired over *m/z* 300-1750 at 70,000 resolution (*m/z* 200) with an AGC target of 1e6, and data-dependent acquisition selected the top 10 most abundant precursor ions for tandem mass spectrometry by HCD fragmentation using an isolation width of 1.6 Da, max fill time of 110ms, and AGC target of 1e5. Peptides were fragmented by a normalized collisional energy of 27, and fragment spectra acquired at a resolution of 17,500 (*m/z* 200).

### 2.4 Proteome of combined sepals and petals (SePel)

Pooled sepals and petals (SePel) of *hot5*-*2* and WT were harvested from the same plants from which pistils were isolated. Material from around 50 flowers at stages before pollination (floral developmental stage, FD 12-13) were pooled and represent one biological replicate (Fig. S4). A total of five biological replicates were further processed as described for the pistil proteome. For the LC-MS\MS run, a 3 µL injection was loaded by a Thermo Easy nLC 1000 UPLC on to a 2 cm trapping column and desalted with 8 µL mobile phase A (0.1% formic acid in water). Peptides were eluted at 300 nL/min on to a 75 µm x 15 cm RSLC column (Thermo) using a linear gradient of 5-35% mobile phase B (0.1% formic acid in acetonitrile) over 90 min. Ions were introduced by positive ESI using a stainless-steel capillary at 2.1 kV into a Thermo Orbitrap Fusion tribrid mass spectrometer. Mass spectra were acquired over m/z 300-1750 at 120,000 resolution (m/z 200) with an AGC target of 1e6, and data-dependent acquisition selected the top 10 most abundant precursor ions for tandem mass spectrometry by HCD fragmentation using an isolation width of 1.6 Da, maximum fill time 110 ms and AGC target 1e5. Peptides were fragmented with a normalized collision energy 27, and fragment spectra acquired at a resolution of 15,000 (m/z 200).

### 2.5 Quantitative proteomics data analysis

Mass spectra for both quantitative proteome datasets (pistil and SePel) were searched against the Uniprot *A. thaliana* database (UP000006548_3702 and UP000006548_3702_additional, downloaded 07/2019) using MaxQuant version 1.6.7.0 (Pistil) or version 1.6.10.43 (SePel) with a 1% FDR at the peptide and protein level, peptides with a minimum length of seven amino acids with carbamidomethylation, N-terminal acetylation and methionine oxidation as fixed modifications. Enzyme specificity was set as C-terminal to arginine and lysine using trypsin as protease and a maximum of two missed cleavages were allowed in the database search. The maximum mass tolerance for precursor and fragment ions was 4.5 ppm and 20 ppm, respectively, with second peptides and match between runs enabled. Label-free quantification was performed with the MaxLFQ algorithm using a minimum ratio count of 2.

The identified protein groups generated by MaxQuant were uploaded to the Perseus program, version 1.6.7.0 (Pistil) or version 1.6.10.50 (SePel)(Tyanova et al., 2016). Site only, reverse, and contaminant peptides were removed from the dataset, LFQ intensities were log2-transformed and missing values imputed assuming a normal distribution. Invalid values were excluded, and empty columns removed. The volcano plot function was used to identify proteins that were significantly changed using a *t*-test with a permutation-based false discovery rate (FDR) of 0.05 and an S0 of 0.1. For hierarchical clustering, LFQ intensities were first z-scored and significantly different proteins (Two sample test, permutation-based FDR of 5%, 250 rounds of randomization) clustered using Euclidean as a distance measure for column and row clustering. 1D annotation enrichment analysis of significant differentially expressed proteins was conducted using a Benjamini-Hochberg FDR of 2 %.

### 2.6 Ubiquitin proteasome activity measurements

Proteolytic activities of the 26S ubiquitin proteasome system (UPS) were measured using fluorogenic substrates following the protocol with modifications described (Üstün and Börnke, 2017). Proteins were extracted from leaves, pistils or inflorescences by resuspending ground powder (in liquid N_2_) of each tissue with extraction buffer (50 mM Hepes-KOH pH 7.7, 0.25 M sucrose, 2 mM ATP, 2 mM DTT) at a 1:1 (w/v) ratio. Samples were centrifuged at 16,000 x g for 10 min at 4 °C to separate cell debris from soluble proteins and the supernatant was subjected to PEG8000 precipitation.

For this, 10 % (v/v) PEG8000 was added to the supernatant and incubated for 30 min at 4 °C. Precipitated proteins were obtained after centrifugation at 16,000 x g for 10 min at 4 °C and the protein pellet resuspended in extraction buffer. Protein concentration was determined using the Bio-Rad assay (Bio-Rad, USA) with BSA as a standard, and concentration adjusted to 1 mg/ml with extraction buffer. For chymotrypsin-like activity measurements 10 µg of protein in assay buffer (100 mM Hepes-KOH pH 7.7, 5 mM MgCl2, 10 mM KCl, 2 mM ATP) was used with 100 µM Suc-LLVY-AMC (Cayman Chemical, USA) as substrate. Trypsin-like and caspase-like activities were determined with 5 µg and 25 µg of protein extract, respectively, and 100 µM of substrate in assay buffer (Ac-RLR-AMC and Z-LLE-AMC for trypsin-like and caspase-like activity, respectively, Cayman Chemical, USA). Samples without protein extract and samples treated with the inhibitor MG132 (80 µM with Suc-LLVY-AMC, 100 µM for Ac-RLR-AMC and Z-LLE-AMC; Sigma, USA) were used as negative controls. Activities were calculated as change in relative fluorescence over time (RFU/min) using a plate reader (BioTek Synergy2, BioTek Instruments, USA).

Fluorescence-based proteasome activity profiling using Me4BodipyFL-Ahx3Leu3VS as substrate was done as described (De Jong et al., 2012), with modifications. Proteins were extracted from leaves, pistils or inflorescences by resuspending ground powder (in liquid N_2_) of each tissue with HR lysis buffer (50 mM Tris-HCl pH 7.5, 0.25 M sucrose, 5 mM MgCl_2_, 2 mM ATP, 1 mM DTT) at a 1:1 (w/v) ratio. For pistil material, 50 pistils per sample were considered one biological replicate. Samples were centrifuged at 16,000 x g for 10 min at 4 °C to separate cell debris from soluble proteins. Protein concentration was measured using the Bio-Rad assay (Bio-Rad, USA) with BSA as standard. To 25 µg of total protein in 25 µl total volume, 5 µM Me4BodipyFL-Ahx3Leu3VS was added and incubated for 1 hr at 30 °C. For negative controls, the UPS inhibitor bortezomib (2 µM) was added to the protein samples and incubated for 30 min at 30 °C prior to substrate labeling. Next, 12.5 µl of 3x reducing sample buffer (120 mM Tris-HCl pH 6.8, 18 % glycerol, 7.5 % 2-mercapto-ethanol, 3 % SDS, 0.006 % bromophenol blue) was added, incubated for 10 min at 70 °C and centrifuged for 1 min at 16 000 xg before loading 10 µl onto 12 % SDS-PAGE gels. After the run, gels were placed in containers with H_2_O and scanned directly using a Typhoon fluorescence scanner (Model FLA9500, GE healthcare, USA) with the following settings: Excitation at 488 nm, Emission at 530 nm, 10 µm pixel size. Band intensities were analyzed with ImageJ (Schneider et al., 2012). After scanning, gels were further washed three times with H_2_O followed by incubation with Coomassie fast stain (Lawrence and Besir, 2009) to visualize total protein.

## 3 Results

### 3.1 Organomercury-resin-assisted capture followed by MS identifies the endogenous S-nitrosoproteome

One of the most severe phenotypes of GSNOR null *hot5*-*2* plants is defective FG development, which results in reduced seed set, illustrating that control of NO-levels is critical for fertility (Lee et al., 2008; Shi et al., 2015; Wang et al., 2024). *hot5-2* plants accumulate RNS, promoting an endogenous increase in protein S-nitrosation (R-SNO), the major biological activity of NO and related RNS (Stamler et al., 1992; Treffon and Vierling, 2022). However, the protein targets of this PTM that may be involved in the *hot5*-*2* fertility defect are unknown. To address the question of which proteins are S-nitrosated in floral tissues of the GSNOR null-mutant *hot5-2*, we utilized a direct enrichment technique for R-SNOs using organomercury (*p*-aminophenylmercuric-acetate) coupled to agarose beads (Doulias and Gould, 2018). The organomercury-resin-assisted capture (MRC) method relies on the specific reaction of organomercury with S-nitrosocysteine, forming a stable thiol-mercury bond (Fig.1). This approach has been used previously in animal systems with no reports published for plants (Doulias et al., 2010; Raju et al., 2012; Doulias et al., 2013). We first performed a pilot experiment using increasing amounts of total protein prepared from leaves of 4- to 6-week-old soil grown *hot5*-*2* plants to test the specificity of the method (Fig. S1). Assaying 5 mg of total protein after alkylation of free thiols already allowed detection of S-nitrosated proteins, and stronger band intensities were obtained with more input material. Importantly, negative controls treated with ascorbate or UV light to remove NO from R-SNOs prior to addition of organomercury resin resulted in little protein recovery, indicating that the method was suitable for further analysis of nitrosated proteins.

**Figure 1.**
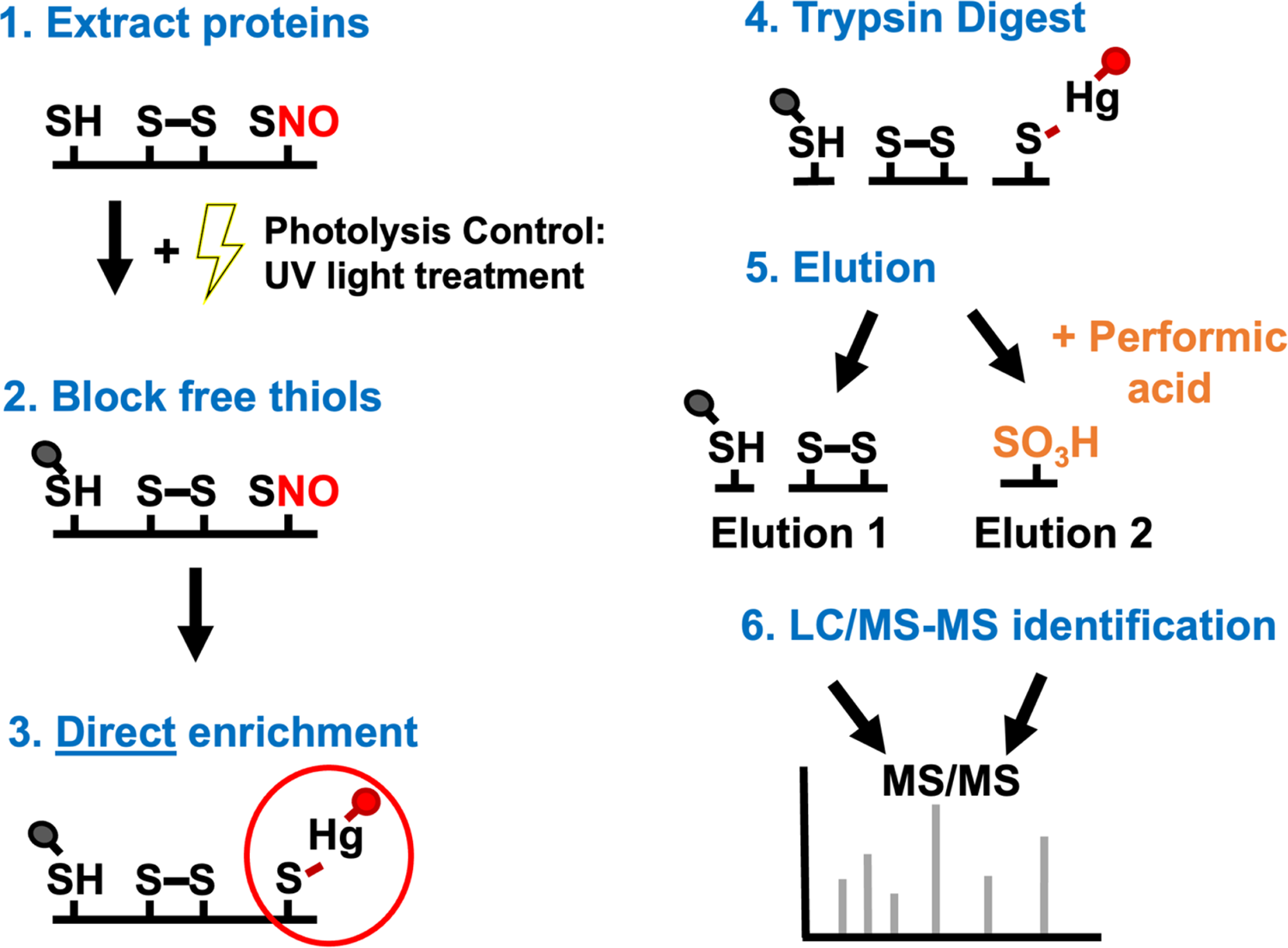
Organomercury-resin-assisted capture (MRC) of S-nitrosated proteins. The MRC method comprises the following steps: (1) Extracting proteins from tissues, (2) blocking reduced cysteines (-SH) with methyl methanethiosulfonate (MMTS), (3) capturing S-nitrosated proteins (-SNO) with *p*-aminophenylmercuric-acetate coupled to agarose beads (Hg), (4) on-column tryptic digestion, (5) release of peptides without (elution 1) or with (elution 2) mild performic acid, and (6) liquid chromatography/tandem MS analysis. UV light treatment (photolysis) of protein extracts removes the NO moiety prior to blocking free thiols and serves as a negative control. Performic acid also oxidizes cysteine thiols to sulfonic acid (-SO_3_H), creating a unique MS signature that permits site-specific identification of the modified cysteines.

### 3.2 *hot5-2* floral tissues have many proteins not previously reported to be S-nitrosated

We utilized the MRC method to assay total protein extracted from entire inflorescences pre-anthesis (corresponding to FD stages up to 12/13) of soil grown WT and *hot5*-*2* plants, with identical samples subjected to UV photolysis as negative controls. We identified a total of 3418 proteins in the set of resin bound proteins in both the experimental and control samples (Fig. 1, elution 1, Materials and Methods). Notably, our S-nitrosated protein dataset included PrxIIE, APX1, and CAT1/3, targets that were identified in other studies (Romero-Puertas et al., 2007; Begara-Morales et al., 2014; Chen et al., 2020; Dreyer et al., 2021), supporting the validity of the MRC method. Using the multiple volcano function (Hawaii plot) in Perseus to filter out proteins identified in the WT, WT photolysis, *hot5*-*2* photolysis samples, we identified 919 S-nitrosated proteins specifically enriched in *hot5-2* floral tissues in elution 1 (Fig. 2A). These proteins grouped into two confidence classes based on false discovery rates (FDRs) of 0.01 (737 protein groups, confidence class A) and 0.05 (an additional 182 protein groups, confidence class B) (Supplemental data 1, Tab 1). Principal component analysis (PCA) confirmed distinct clustering of *hot5*-*2* compared to WT, WT photolysis and *hot5*-*2* photolysis controls (Fig. 2B).

**Figure 2.**
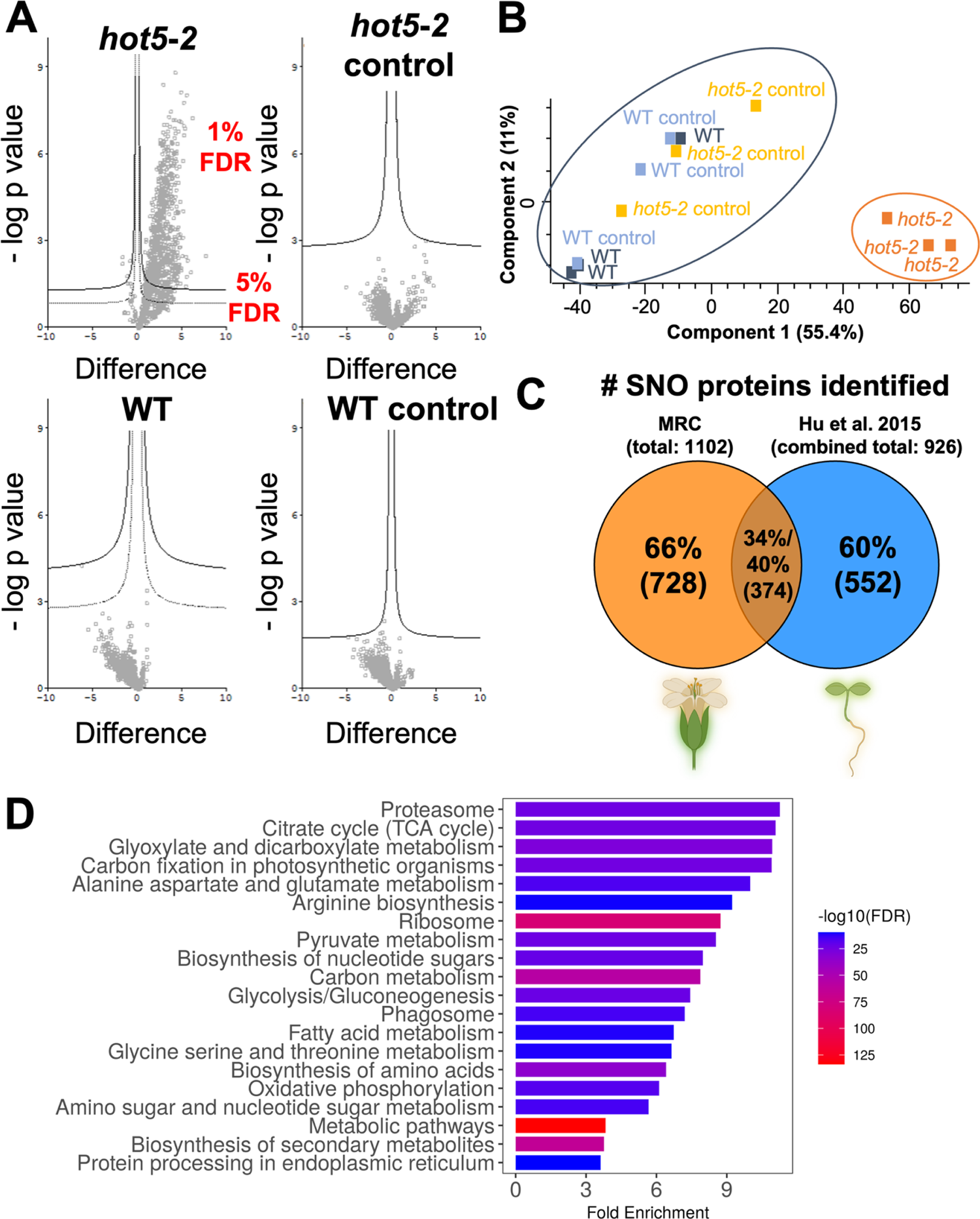
The endogenous S-nitrosoproteome of *hot5-2* floral tissues. (A) Hawaii-plot (multiple volcano plot) of S-nitrosated proteins from WT and *hot5-2* inflorescences pre-anthesis. UV photolysis of *hot5-2* or WT represent controls. The significance (-log p value) is plotted against the t-test difference comparing all other samples. The two cut-off lines define two confidence classes with S0 = 0.1 and FDRs of 0.01 (solid line) and 0.05 (dashed line). Identified proteins are listed in the Supplemental data set 1, Tab 1 and 2. All samples were prepared in triplicates. (B) Principal component analysis (PCA) plot of identified S-nitrosated proteins indicates differences between WT, WT control, *hot5-2* control compared to *hot5*-*2* samples. (C) Comparison of S-nitrosated proteins identified in *hot5-2* from 15-day-old seedlings (Hu et al., 2015) versus inflorescences in this study. (D) KEGG enrichment analysis of S-nitrosated proteins using ShinyGo. The top 20 most enriched pathways with false discovery rate (-log10 FDR) are shown. A list of proteins associated with the different terms is provided in Supplemental Data Set 3.

One advantage of the MRC method is that it can provide site-specific information on Cys residues that are modified, because when resin-bound R-SNO-peptides are oxidatively released by performic acid R-SO_3_H modified peptides are produced (Fig. 1, elution 2, see Materials and Methods). A total of 533 S-nitrosated peptides representing 183 protein groups were identified in at least one of the three biological replicates in the performic acid elutions of WT and/or *hot5-2*, excluding peptides that appeared only in the photolysis controls (Supplemental data 1, Tab 3). Of these 533 R-SNO peptides 359 (67%; 295 protein groups) and 48 (9%; 39 protein groups) were unique to *hot5-2* and WT, respectively, while the other 126 (24%; 103 protein groups) were identified in both genotypes (Supplemental data 1, Tab 4 and 5). Surprisingly, of the 533 R-SNO peptides, only 303 (57%) matched the 919 S-nitrosated proteins identified in *hot5-2*. Therefore, in total we found 1049 S-nitrosated protein groups unique to *hot5-2*, when considering both the peptides from elution 1 and the R-SO_3_H modified peptides from elution 2 (Supplemental data 1, Tabs 1 and 4).

We further compared our total data set of 1102 S-nitrosoproteins (919 elution 1; 183 elution 2) from *A. thaliana* floral tissues (1049 specific to *hot5-2*) to previous studies examining the nitrosoproteome in *A. thaliana*. Hu et al. (2015) identified 926 nitrosated proteins in 15 day-old *hot5*-*2* mutant and WT seedlings (Hu et al., 2015) (Fig. 2C and Supplemental data 1, Tab 8-10). 374 proteins (∼34%) were identified in both our inflorescence and the Hu et al. (2015) seedling experiments, whereas 728 (∼66%) and 552 proteins (∼60%) were unique to the inflorescence or seedling data sets, respectively (Fig. 2C and Supplement data 1, Tab 2 and 7). Comparing specifically R-SNO peptides, 131 were identified in both studies, of which only 58 (44%) are the same peptides, whereas 73 (56%) are different, suggesting that the same proteins can be S-nitrosated at different Cys residues. We also compared our dataset to S-nitrosated proteins identified in GSNO-treated *A. thaliana* cell culture extracts and NO-treated leaves (Lindermayr et al., 2005). Our data included 31 proteins of 63 nitrosated proteins identified in cell culture extracts and 36 of 52 proteins identified in NO-treated leaves (Supplemental data 1, Tab 7, 8, 11, 12). Taken together, these data suggest that there are likely differences in S-nitrosated proteins depending on plant developmental stage, as well as differences between effects of elevated endogenous NO in *hot5-2* plants, versus exogenously applied NO donors.

### 3.3 S-nitrosated proteins in floral tissues are implicated in multiple metabolic pathways

To gain insight into which metabolic and regulatory pathways may be impacted by protein S-nitrosation in inflorescences, we classified the 1102 S-nitrosated protein groups we identified using MapMan (Usadel et al., 2009) (Fig. S2A, B). S-nitrosated proteins mapped to multiple pathways, including photosynthesis and the Calvin cycle (45 proteins), glycolysis (32 proteins) and the TCA cycle (36 proteins) (Supplement data 2). We also identified 30 S-nitrosated proteins involved in transcript regulation (e.g. AGO4), and several redox-related proteins, including glutathione peroxidases (GPX2/5/6), ascorbate peroxidases (APX1/3), monodehydroascorbate reductases (MDAR1/2) and dehydroascorbate reductase (DHAR1). Seventy-seven proteins were classified in protein degradation pathways.

KEGG pathway analysis of the total data set of 1102 S-nitrosated proteins using ShinyGo (Ge et al., 2020) complemented the MapMan classification showing enrichment in proteins involved in carbon fixation (Fig. 3A), the TCA cycle (Fig. 3B) and oxidative phosphorylation (Fig. 4A). Of the 37 identified S-nitrosated proteins that correspond to the KEGG term oxidative phosphorylation, 65 % (24 proteins) are newly identified and specific for floral tissues. Significantly, 31 *hot5-2* specific S-nitrosated proteins are also associated with the KEGG term proteasome, of which 15 and 10 represent components of the regulatory or core subunits of the 26S ubiquitin proteasome system, respectively (Fig. 2D, Fig. 4B and Supplemental data 3). Of those 31 proteasomal proteins, 55% (17 proteins) are components of the 26S proteasome identified in floral tissues and that have not been reported previously as targets for S-nitrosation (Fig. 4B).

**Figure 3.**
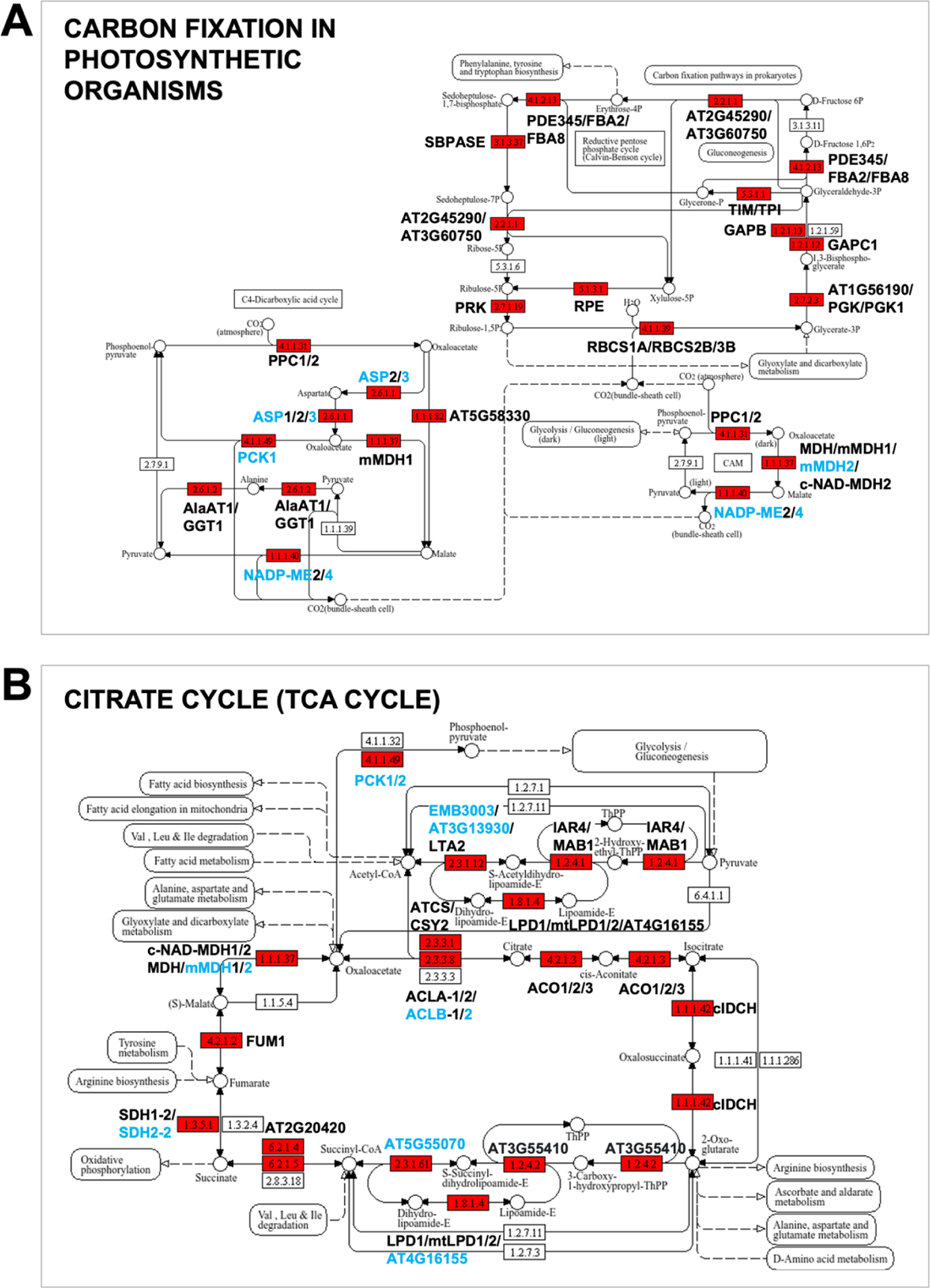
KEGG pathway diagram highlighting enriched S-nitrosated proteins involved in carbon fixation (A) and TCA cycle (B). Shown in red are the EC code numbers of identified S-nitrosated proteins and highlighted in blue are the S-nitrosated proteins identified in inflorescences only. Proteins associated with the different terms are listed in Supplemental Data Set 3. Abbreviations in **(A):** AlaAT1, AT1G17290; GGT1, AT1G23310; PRK, AT1G32060; GAPB, AT1G42970; mMDH1, AT1G53240; PPC1, AT1G53310; AT1G56190; RBCS1A, AT1G67090; PGK, AT1G79550; NADP-ME4, AT1G79750; PDE345, AT2G01140; TIM, AT2G21170; ASP1, AT2G30970; PPC2, AT2G42600; AT2G45290; GAPC1, AT3G04120; PGK1, AT1G79550; mMDH2, AT3G15020; MDH, AT3G47520; FBA8, AT3G52930; TPI, AT3G55440; SBPASE, AT3G55800; AT3G60750; PCK1, AT4G37870; FBA2, AT4G38970; ASP3, AT5G11520; NADP-ME2, AT5G11670; ASP2, AT5G19550; RBCS3B, AT5G38410; RBCS2B, AT5G38420; c-NAD-MDH2, AT5G43330; AT5G58330; RPE, AT5G61410. Abbreviations in **(B)**: c-NAD-MDH1, AT1G04410; ACLA-1, AT1G10670; IAR4, AT1G24180; EMB3003, AT1G34430; mtLPD1, AT1G48030; mMDH1, AT1G53240; ACLA-2, AT1G60810; cICDH, AT1G65930; ACO3, AT2G05710; SDH1-2, AT2G18450; AT2G20420; ATCS, AT2G44350; FUM1, AT2G47510; ACLB-1, AT3G06650; AT3G13930; mMDH2, AT3G15020; LPD1, AT3G16950; mtLPD2, AT3G17240; LTA2, AT3G25860; MDH, AT3G47520; AT3G55410; CSY2, AT3G58750; AT4G16155; ACO2, AT4G26970; ACO1, AT4G35830; PCK1, AT4G37870; SDH2-2, AT5G40650; c-NAD-MDH2, AT5G43330; ACLB-2, AT5G49460; MAB1, AT5G50850; AT5G55070; PCK2, AT5G65690.

**Figure 4.**
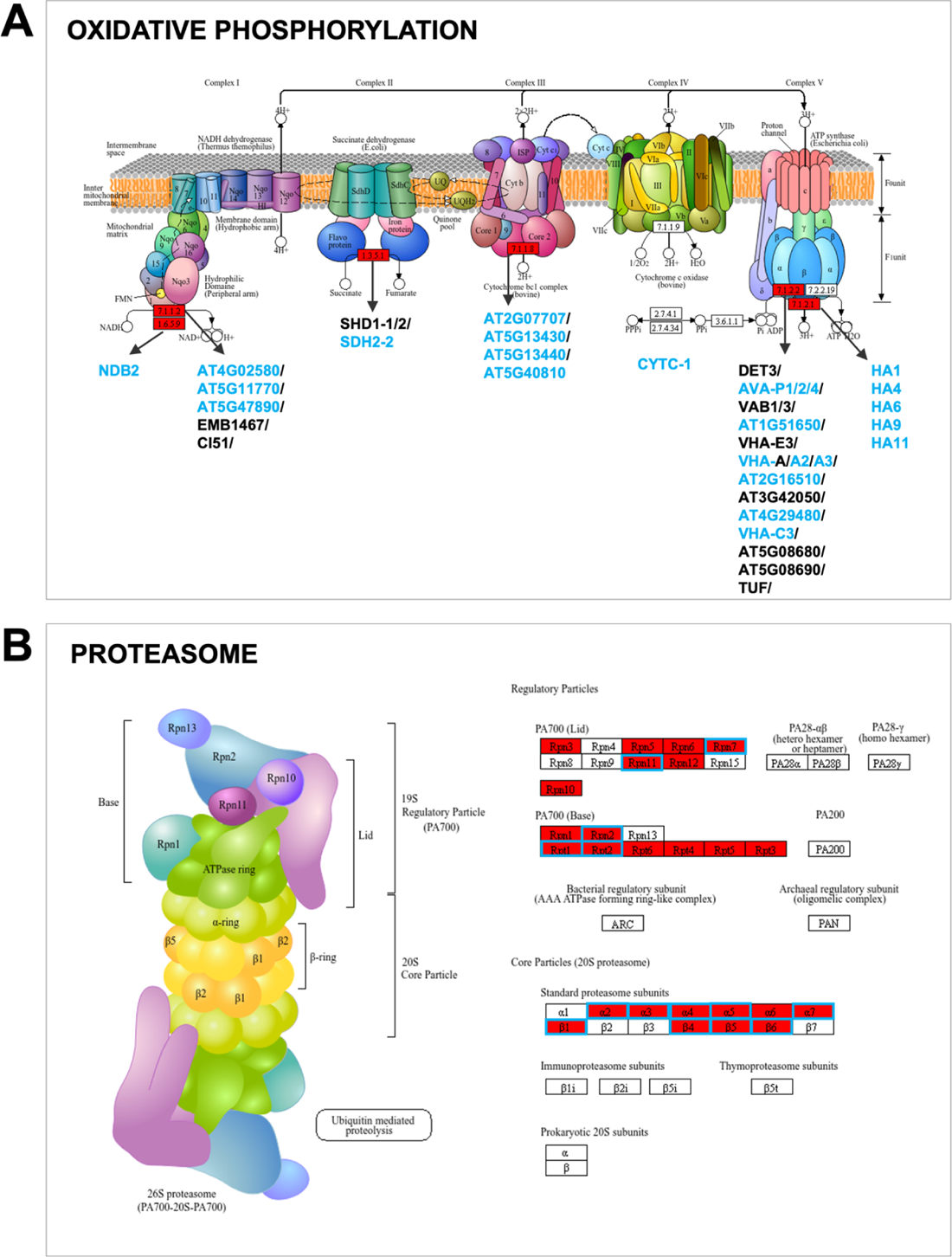
KEGG pathway diagram highlighting enriched S-nitrosated proteins involved in oxidative phosphorylation (A) and protein degradation (B). Shown in red are the EC code numbers of identified S-nitrosated proteins and highlighted in blue are the S-nitrosated proteins identified in inflorescences only. Proteins associated with the different terms are listed in Supplemental Data Set 3. Abbreviations in **(A)**: DET3, AT1G12840; AVA-P2, AT1G19910; VAB3, AT1G20260; CYTC-1, AT1G22840; AT1G51650; VHA-E3, AT1G64200; AVA-P4, AT1G75630; VAB1, AT1G76030; VHA-A, AT1G78900; HA9, AT1G80660; HA6, AT2G07560; AT2G07707; AT2G16510; SDH1-2, AT2G18450; HA1, AT2G18960; VHA-A2, AT2G21410; AT3G42050; HA4, AT3G47950; AT4G02580; NDB2, AT4G05020; TUF, AT4G11150; AT4G29480; AVA-P1, AT4G34720; VHA-C3, AT4G38920; VHA-A3, AT4G39080; CI51, AT5G08530; AT5G08680; AT5G08690; AT5G11770; AT5G13430; AT5G13440; EMB1467, AT5G37510; SDH2-2, AT5G40650; AT5G40810; AT5G47890; HA11, AT5G62670; SDH1-1, AT5G66760. Abbreviations in **(B)**: Rpn1/RPN1A, AT2G20580; Rpn2/RPN2A, AT2G32730; Rpn3/EMB2719, AT1G20200; Rpn5/EMB2107, AT5G09900; Rpn6/ATS9, AT1G29150; Rpn7/RPN7, AT4G24820; Rpn10/RPN10, AT4G38630; Rpn11/RPN11, AT5G23540; Rpn12/RPN12a, AT1G64520; Rpt1/RPT1A, AT1G53750; Rpt2/RPT2a, AT4G29040; Rpt2/RPT2b, AT2G20140; Rpt3/RPT3, AT5G58290; Rpt4/RPT4A, AT5G43010; Rpt5/RPT5A, AT3G05530; Rpt5/RPT5B, AT1G09100; Rpt6/RPT6A, AT5G19990; Rpt6/RPT6B, AT5G20000; α2/PAB1, AT1G16470; α2/ PAB2,AT1G79210; α3/PAC1, AT3G22110; α4/PAD1, AT3G51260; α5/PAE1, AT1G53850; α5/PAE2, AT1G53850; α6/PAF1, AT5G42790; α6/PAF2, AT1G47250; α7/PAG1, AT2G27020; β1/PBA1, AT4G31300; β4/PBD1, AT3G22630; β5/PBE1, AT1G13060; β6/PBF1, AT3G60820.

Notably, KEGG pathway analysis of the 728 inflorescence-specific S-nitrosated protein groups (Fig. 2C) revealed a similar enrichment pattern of photosynthesis- and proteasomal-related processes as shown for the entire dataset. In addition, KEGG-terms associated with “ribosome” and “fatty acid metabolism” are enriched in *hot5-2* inflorescences (Supplemental data 3).

### 3.4 Comparison of the quantitative proteome with the S-nitrosoproteome of *hot5-2* floral tissues

The specific enrichment of S-nitrosated proteins in floral tissues of *hot5-2* compared to WT could partially result from an upregulation in protein abundance. In addition, recent data demonstrated that the *hot5-2* fertility defect is mainly a result of defects in female gametophyte development (Wang et al., 2024). It was, therefore, of interest to analyze the total quantitative proteome of floral tissues to address the relationship of protein nitrosation and protein levels, as well as to understand how differences in protein levels might contribute to *hot5-2* fertility defects. While the amount of material required for the MRC analysis limited our experiments to combined floral tissues, for total quantitative proteomics we were able to analyze isolated pistils compared to a sample of pooled sepals and petals (SePels) of *hot5-2* and WT at flower developmental stages 12-13 prefertilization (Fig. S4). We identified a total of 3297 and 3353 proteins in pistils and SePels, respectively (Fig. 5A and B), of which approximately only 6 % (193 proteins; pistil) and 4 % (149 proteins; SePel) were differentially regulated in *hot5*-*2* compared to the WT control (Supplement data 4 and 5).

**Figure 5.**
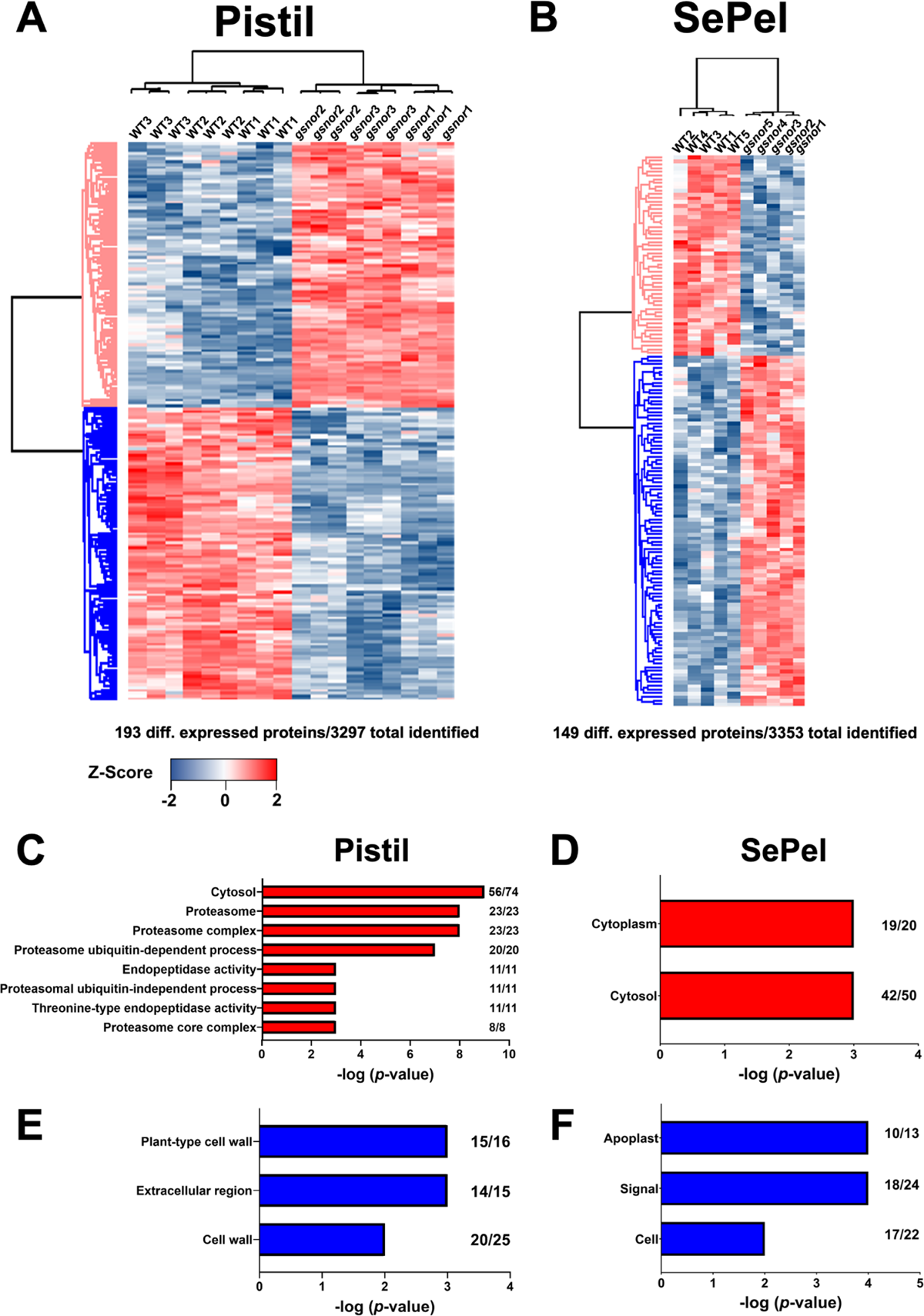
LFQ proteomic analysis of floral tissues of WT and *hot5-2* and Gene ontology (GO) enrichment of differentially regulated proteins. Heatmap of differentially expressed proteins in WT and *hot5*-*2* pistils (A) and (B) combined sepals and petals sample (SePel). Three biological replicates with three technical replicates for pistils and five biological replicates for SePel were analyzed. LFQ intensities were Z-scored prior to Euclidean distance-based hierarchical clustering with Perseus. Blue and red indicate a lower and higher abundance for each protein in the samples. Up-(C, D; red) and downregulated (E, F; blue) clusters of pistils and SePels, respectively. *p*-values of significant GO-terms regarding biological processes, molecular functions and cellular compartment were –log transformed and displayed in bar graphs. Numbers at the end of each bar represent the number of proteins in that cluster that are associated with the specific GO term. A list of the differentially regulated proteins is provided in Supplemental Data Set 4 and 5.

These significantly up- or down-regulated proteins for each tissue were further clustered and gene set enrichment analysis revealed an enrichment of proteasomal terms in upregulated proteins (cluster 1) in *hot5-2* pistils (Fig. 5C), whereas down-regulated proteins in cluster 2 included processes related to the extracellular region (Fig. 5E). In comparison to pistils, differentially regulated proteins in SePels were associated with fewer GO-terms, which were dominated by “cytoplasm” for up- and “apoplast”, “signal” and “cell” for down-regulated proteins (Fig. 5 D and F). These data clearly demonstrate that the overall protein composition is altered in different floral tissues in *hot5*-*2* compared to WT.

For further analysis, as described below, we included our previously published quantitative proteome dataset (Supplemental data 6) obtained from leaves of 4-6 week old, soil grown *hot5-2* and WT plants (Treffon et al., 2021) as an additional control for differences in these proteomes. We reasoned that by comparing the protein expression levels of different protein groups among all three samples (pistils, SePel, leaves) that we could identify proteins that are exclusively enriched or depleted in a specific tissue as well as overall differences between *hot5-2* and WT.

### 3.5 UDP-glycosyltransferases (UGTs) show specific regulation in *hot5-2*

We evaluated the most highly up-or downregulated proteins in the three different tissues using volcano plot analysis (Fig.6). One of the most up-regulated protein groups in *hot5-2* pistils belongs to the UDP-glycosyltransferase protein family (UGT). Of the 17 UGTs identified in pistils, seven were significantly upregulated (Fig. 6A and Table1). UGTs are involved in multiple pathways in plants. They act to glycosylate various substrates and are reported to be important for modifying plant hormones including auxins, salicylic acid and abscisic acid thereby regulating hormone bioavailability (Dean and Delaney, 2008; Tognetti et al., 2010; Bock, 2016; Trujillo-Hernandez et al., 2020). In addition, UGT75B1 has been identified as a component of the callose synthase complex involved in callose deposition at the forming cell plate during cytokinesis (Hong et al., 2001). The SePel and leaf samples also showed significant changes in certain UGTs, but in a different pattern of regulation as well as in which UGTs were detected in the three tissues (Fig. 6A and Table 1). Overall, UGTs are responding very differently to the disturbed NO homeostasis throughout *hot5-2* mutant plants, which may prove significance relative to the observed fertility defects (Lee et al., 2008; Wang et al., 2024).

**Figure 6.**
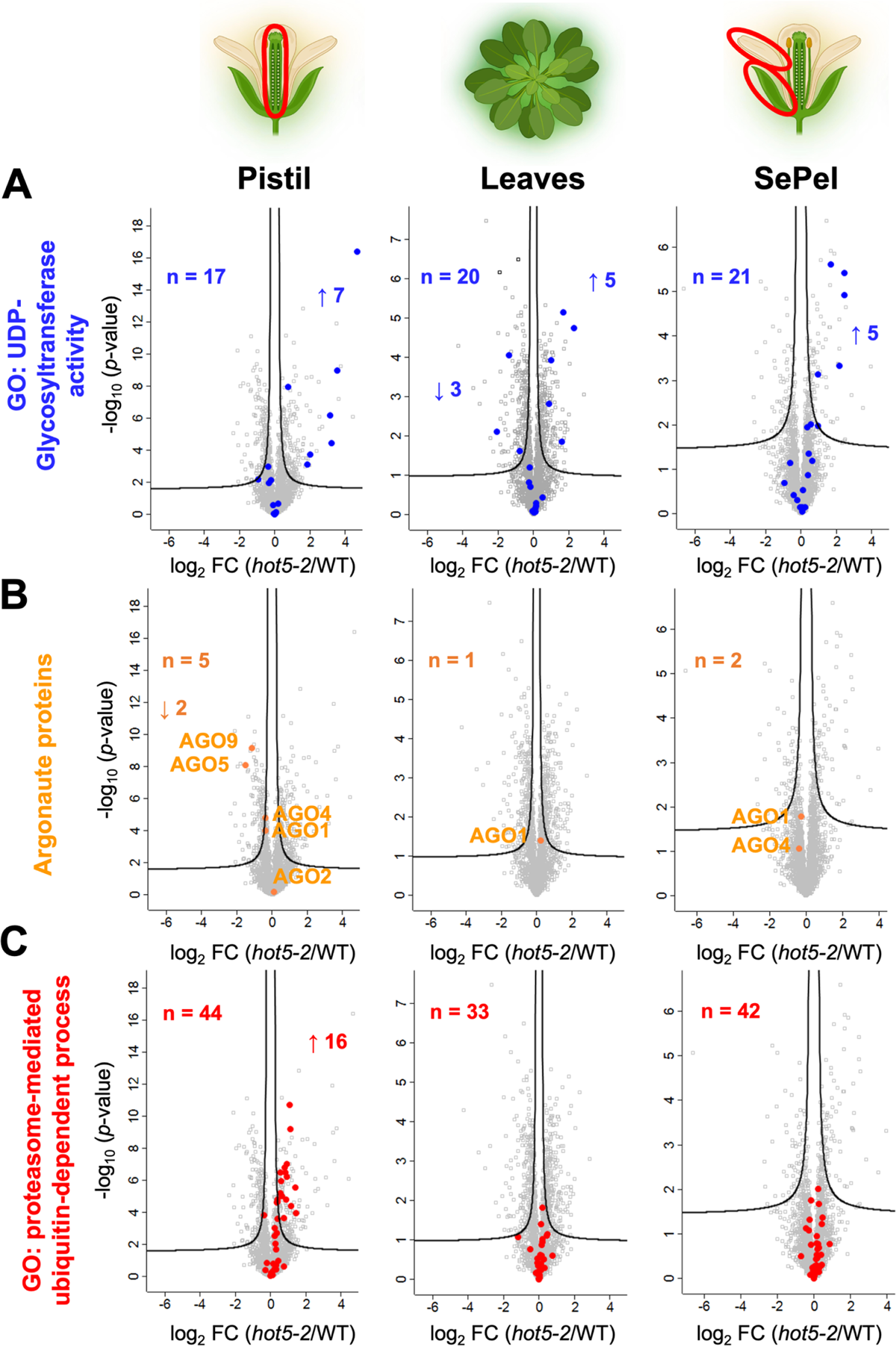
Volcano plots of differentially regulated proteins in different *hot5-2* tissues. The pistil and SePel dataset are reported in this study, whereas the leaf dataset was published previously (Treffon et al., 2021). Proteins associated with the 26S proteasome (**A**) GO-term: proteasome-mediated ubiquitin-dependent process), UDP-glycosyltransferases (**B**) GO-term: UDP-glycosyltransferase activity) and Argonaute proteins **(C)** are illustrated. The number of detected up- and down-regulated proteins for each group and the total number of identified proteins in the respective dataset are shown. The -log10 of the corrected *p*-values is plotted against the log2-fold change (FC) in protein levels. FDR cutoff (5%; S0= 0.1) is indicated by black lines. A list of the significantly regulated proteins for each tissue is provided in the Supplemental Data Sets 4-6.

**Table 1.**
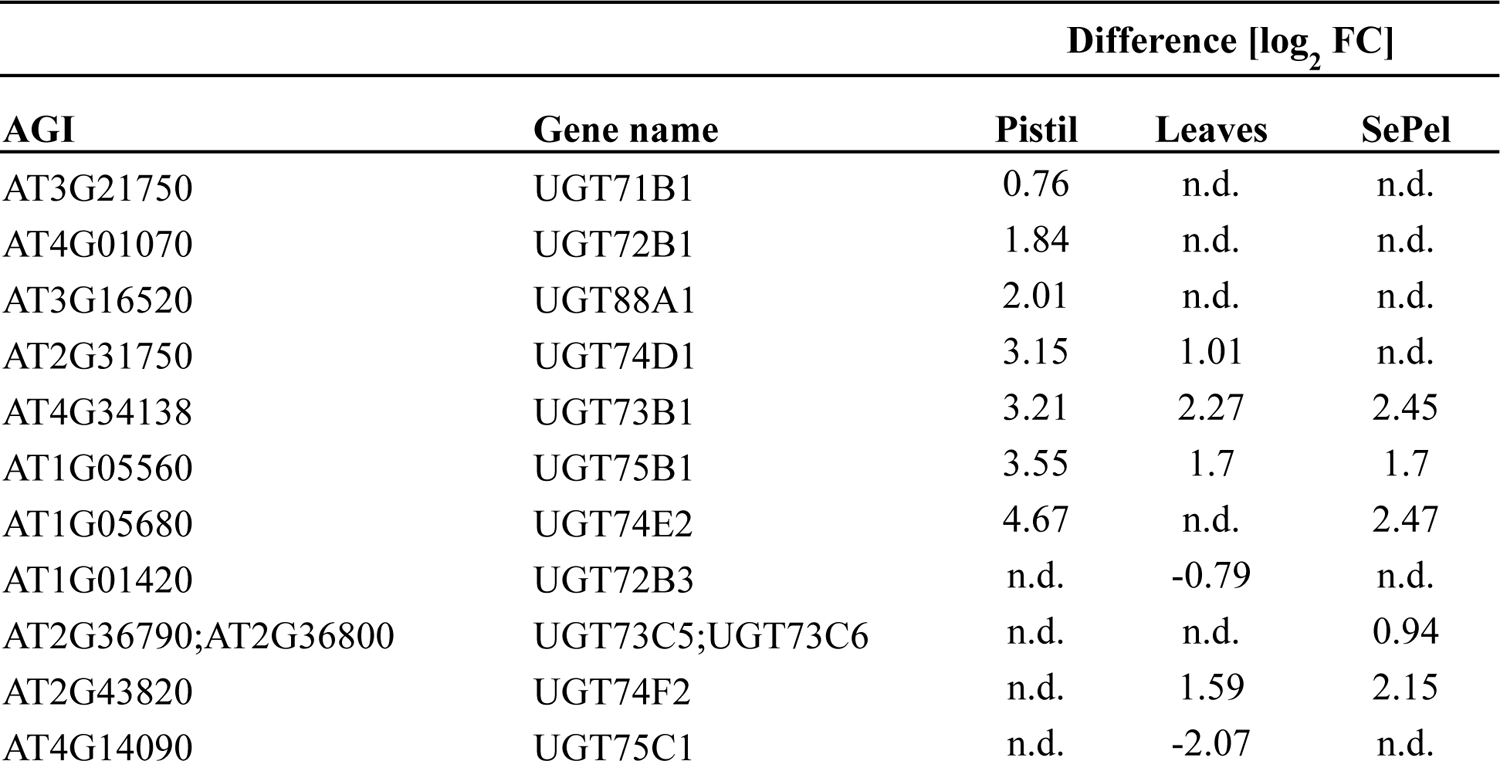
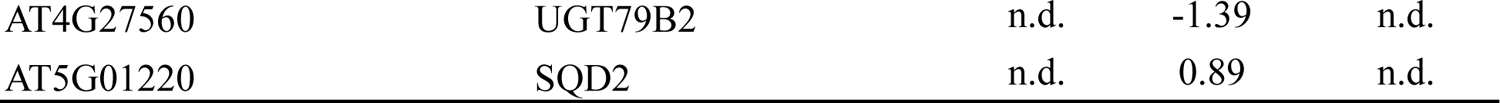
Differentially regulated UGT proteins in *hot5-2*. List of significantly up- or downregulated UGT proteins identified in pistils, leaves or SePels with log_2_ fold change values and ranked based on FC in pistils. Proteins not identified in a specific sample are labeled with “n.d”.

| Difference [log<sub>2</sub> FC] |  |  |  |  |
| --- | --- | --- | --- | --- |
| AGI | Gene name | Pistil | Leaves | SePel |
| AT3G21750 | UGT71B1 | 0.76 | n.d. | n.d. |
| AT4G01070 | UGT72B1 | 1.84 | n.d. | n.d. |
| AT3G16520 | UGT88A1 | 2.01 | n.d. | n.d. |
| AT2G31750 | UGT74D1 | 3.15 | 1.01 | n.d. |
| AT4G34138 | UGT73B1 | 3.21 | 2.27 | 2.45 |
| AT1G05560 | UGT75B1 | 3.55 | 1.7 | 1.7 |
| AT1G05680 | UGT74E2 | 4.67 | n.d. | 2.47 |
| AT1G01420 | UGT72B3 | n.d. | -0.79 | n.d. |
| AT2G36790;AT2G36800 | UGT73C5;UGT73C6 | n.d. | n.d. | 0.94 |
| AT2G43820 | UGT74F2 | n.d. | 1.59 | 2.15 |
| AT4G14090 | UGT75C1 | n.d. | -2.07 | n.d. |
| AT4G27560 | UGT79B2 | n.d. | -1.39 | n.d. |
| AT5G01220 | SQD2 | n.d. | 0.89 | n.d. |

We captured four UGT protein groups in our S-nitrosoproteome dataset (UGT79B2/B3, UGT78D1, UGT89C1, UGT80A2), but these were not found to be up- or down-regulated (Supplement data 7). Nevertheless, the R-SNO analysis of *A. thaliana hot5-2* seedlings by Hu et al. (2015) reported that 9 of the 17 upregulated UGTs in our datasets were S-nitrosated. The absence of these UGTs from our S-nitrosoproteome dataset could reflect tissue specific differences in S-nitrosation, although we cannot rule out differences in the methods of SNO capture.

### 3.6 Specific argonaute (AGO) proteins are down-regulated in *hot5-2* pistils

In contrast to the UGTs, two argonaute proteins, AGO5 and AGO9, are significantly downregulated in *hot5-2* pistils compared to the other tissues (Fig. 6B). In *A. thaliana*, as in many other organisms, AGO proteins are integral components of the RNA-induced silencing complex (RISC), playing crucial roles in RNA interference and post-transcriptional gene regulation (Meister, 2013; Fang and Qi, 2016). *A. thaliana* has a relatively small family of ten AGO proteins in three clades: AGO1-like, AGO2-like and AGO4-like (Hernandez-Lagana et al., 2016). Notably, AGO5 and AGO9 (AGO1-like and AGO4-like, respectively) are involved in female reproductive development. AGO5 mutants show defects in the initiation of megagametogenesis (Tucker et al., 2012), while AGO9 acts in nucellar epidermal cells inhibiting the formation of multiple megaspore-like cells during early ovule development (Olmedo-Monfil et al., 2010). Although AGO5 and AGO9 were not found in our S-nitrosoproteome data, AGO1 and AGO4 were identified as S-nitrosated (Supplement data 1). We obtained five SNO-peptides for AGO4, which was reported recently to be S-nitrosated and important for the NO-dependent regulation of shoot meristem activity through its impact on DNA methylation (Zeng et al., 2023). Taken together, our results indicate that NO is a regulator of expression and posttranslational modification of multiple AGO proteins, of which two function during early female development.

### 3.7 Class 4 Aldo-keto reductases are upregulated and S-nitrosated in *hot5-2*

While GSNOR is clearly a major regulator of NO homeostasis in all organisms, aldo-keto reductases (AKRs) have been recently described in both mammals (Stomberski et al., 2019) and plants (Treffon et al., 2021) as additional enzymes capable of regulating NO homeostasis by catabolizing GSNO. While leaves of *hot5-2* show an upregulation of two of four AKRs in the subclade 4C (AKR4C8 and AKR4C9) (Treffon et al., 2021), pistils in *hot5-2* also accumulate AKR4C10 and C11, suggesting that those two proteins are mainly expressed in floral tissues (Fig. S3). We also found that AKR4C8 and 9 are targets for S-nitrosation in *hot5-2* inflorescences (Supplemental data 1, Tab 1). Further biochemical studies and genetic analysis with plant mutants are needed to assess the significance of AKR4C proteins in regulating GSNO levels at different physiological and developmental stages.

### 3.8 Major changes in abundance of proteasomal components in relation to S-nitrosation

In parallel to the overall enrichment of 26S proteasomal-related GO-terms (Fig. 2D) together with an increased number of S-nitrosated UPS proteins in floral tissues (Fig. 4B, Supplement data 1 and 3), we observed a significant up-regulation in protein abundance of UPS proteins in pistils of *hot5-2* (16 out of a total of 44 UPS proteins detected, Fig. 6C). Proteins with log2-fold changes of 0.56-1.45 included seven regulatory particle proteins (RPN2B, RPN3A, RPN5A, RPN7, RPN8B, RPN9B, RPN12A) and seven core proteins (PBD2, PAB2, PAD1, PAE1, PAA2, PAD2, PAE2), as well as proteins reported to be involved in proteasome activation (PA200) and assembly (At3g15180) based on the UniProt database (Table 2). Notably, we recovered 10 of these 16 upregulated 26S proteasomal proteins in our S-nitrosoproteome experiment (Table 2 and Fig. S5). We identified one S-nitrosopeptide for RPN5A (IDRPSGIVC(397)FQIAK), which differs from the RPN5A nitrosated peptide reported by Hu et al. (2015), indicating that RPN5A can be S-nitrosated at multiple positions. Furthermore, we identified 11 additional R-SNO peptides from 11 proteins in elution 2 (Table 2). Importantly, although approximately the same number of UPS proteins were identified in the leaves and SePel data sets (33 and 42, respectively, Fig. 6C), no abundance changes were detected in these tissues in *hot5-2* compared to WT, suggesting that UPS proteins are exclusively upregulated and S-nitrosated in *hot5-2* pistils.

**Table 2.**
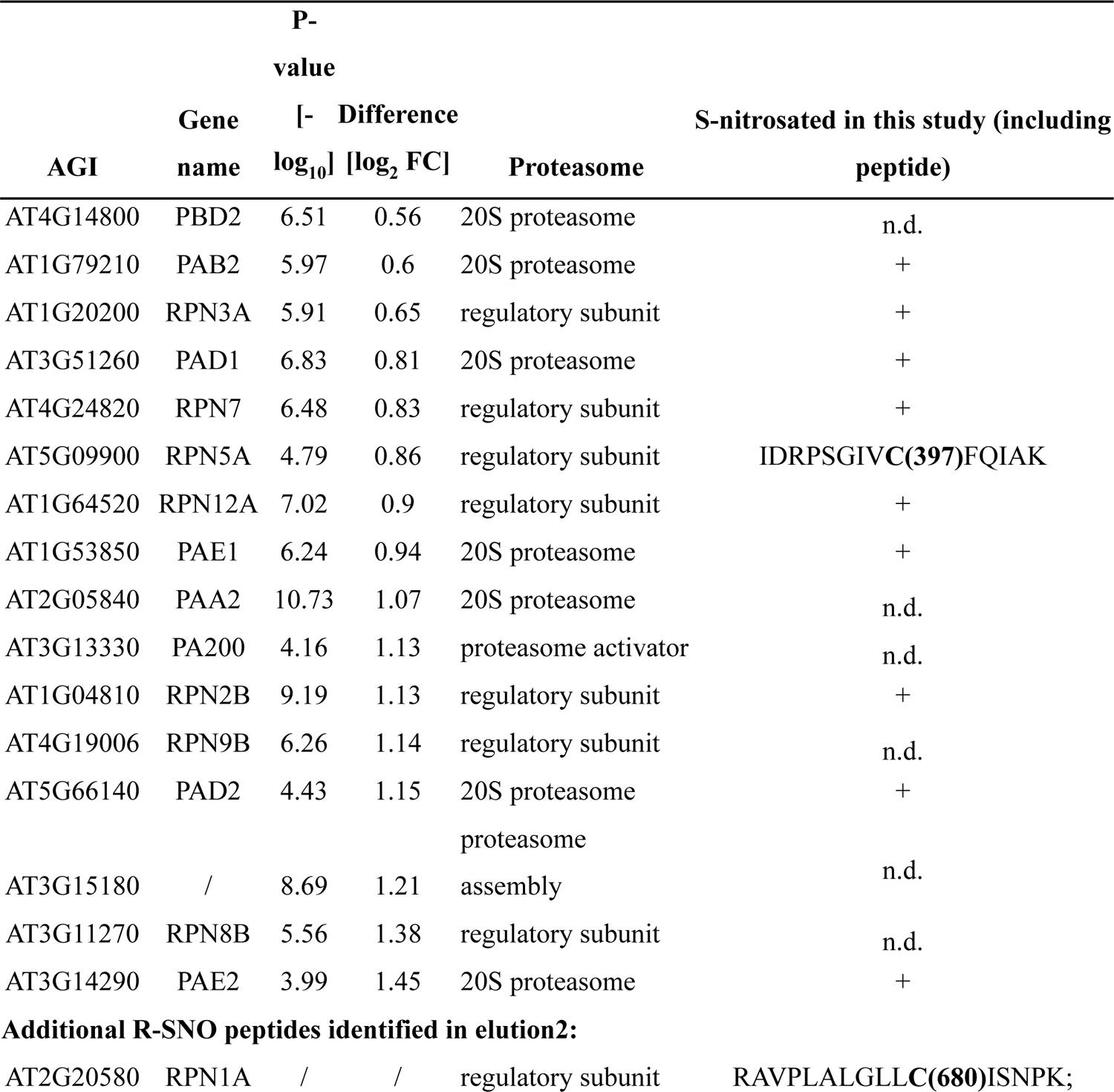
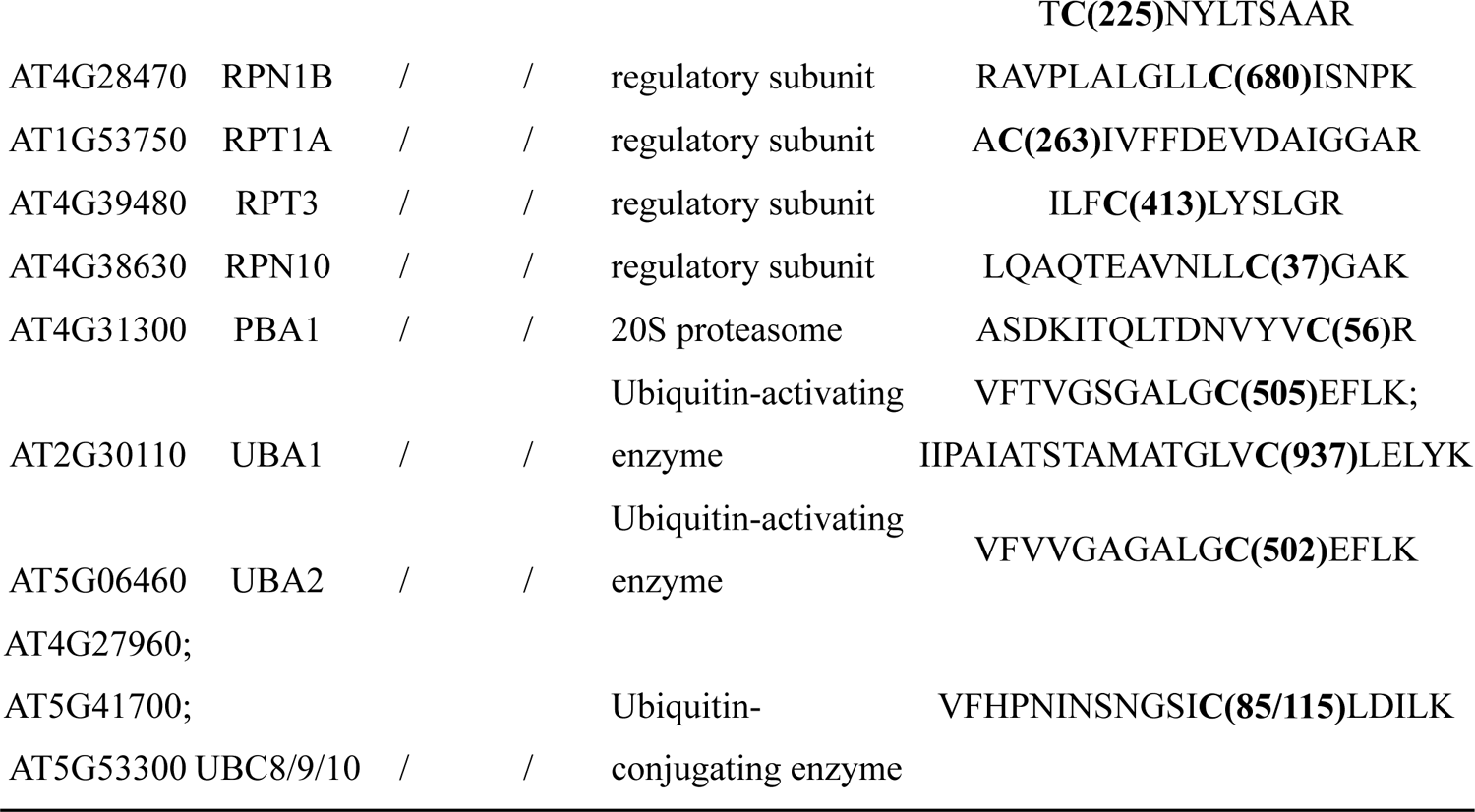
Upregulated and S-nitrosated UPS proteins in floral tissues of *hot5-2*. List of differentially regulated and S-nitrosated UPS proteins with AGI number and calculated *p*-value (-log_10_) that are significantly enriched and sorted by log_2_-fold-change. See also Figure S5. Identified S-nitrosated proteins are marked with a “+”, whereas differentially regulated proteins not captured in our SNO-proteomics experiments are labeled with “n.d.”. RPN5A and RPN12A were also reported in Hu et. al. (2015).

To explore further any potential correlation between changes in protein abundance and the detection of S-nitrosated proteins in different *hot5-2* tissues, we compared the datasets based on the log2-fold change values of significantly up- or down-regulated proteins and our R-SNO data set (elution 1) with each other. Pearson correlation coefficients for up- or downregulated proteins from pistils (0.15 and 0.27), leaves (0.17 and 0.23) and SePels (0.46 and 0.46) in comparison to the S-nitrosoproteome reveal a low correlation in general (Supplemental data 8). However, out of the 497 differentially regulated proteins in *hot5-2* leaves, 27 % (134 proteins) were also identified in our S-nitrosoproteome dataset, with 46 % (62 proteins) and 54 % (72 proteins) found to be up- and down regulated, respectively (Supplemental data 8). In contrast, we found that a higher number of significantly upregulated proteins in pistils (75 %) and SePels (70 %) are S-nitrosated, indicating that there is a positive correlation between protein abundance and S-nitrosation status in *hot5-2* floral tissues.

### 3.9 Trypsin-like activity of the 26S proteasome is specifically increased in pistils of *hot5-2*

Considering the upregulation of 26S proteasomal proteins in pistils, as well as an increased number of S-nitrosated UPS proteins in *hot5-2*, we investigated the different UPS protease activities in leaf, pistil, and inflorescence material of WT and GSNOR mutant plants towards the different UPS protease activities using fluorescently labeled substrates (Fig. 7). The main component of the UPS, the 20S core, shows distinct chymotrypsin-, trypsin- and caspase-like activities (Marshall and Vierstra, 2019). Using standard fluorescently labeled substrates (see Materials & Methods), we assayed protease activities in tissue extracts. We did not detect differences in chymotrypsin- or caspase-like activities when comparing different *hot5-2* tissues with WT (Fig. 7A, C), and trypsin-like activity showed no alteration in *hot5-2* leaves and inflorescences compared to WT (Fig. 7B). However, *hot5-2* pistils exhibited a 42% increased trypsin-like proteolytic activity relative to the WT pistil sample (Fig. 7B). We further assayed proteasome proteolytic activity by gel electrophoresis using the fluorescent probe Me4BodipyFL-Ahx3Leu3VS, which has allowed for the identification and activity determination of specific proteasome core subunits in mammals (De Jong et al., 2012; Leestemaker et al., 2017). Me4BodipyFL-Ahx3Leu3VS comprises a fluorescent tag and a proteasome-targeting motif with a vinyl sulfone group that covalently reacts with the N-terminal threonine of all active core subunits (De Jong et al., 2012). Multiple bands at around 25 kDa were detected, of which most were absent when extracts were treated with the proteasome inhibitor bortezomib, indicating that the signals result from proteins with endopeptidase activity (Fig. 7D). We cannot assign individual bands to specific proteasome core proteins, but quantification of all the bands indicate that floral tissues of both *hot5-2* and WT showed an increased proteasomal activity when compared to leaves (Fig. 7E). In addition, there is an 81 % increase in fluorescence signal for *hot5-2* pistils compared to the WT pistils sample. Combined with our proteolytic activity assays, these data suggest that a deregulated NO homeostasis results in an increased trypsin-like UPS activity in pistils of *hot5-2*.

**Figure 7.**
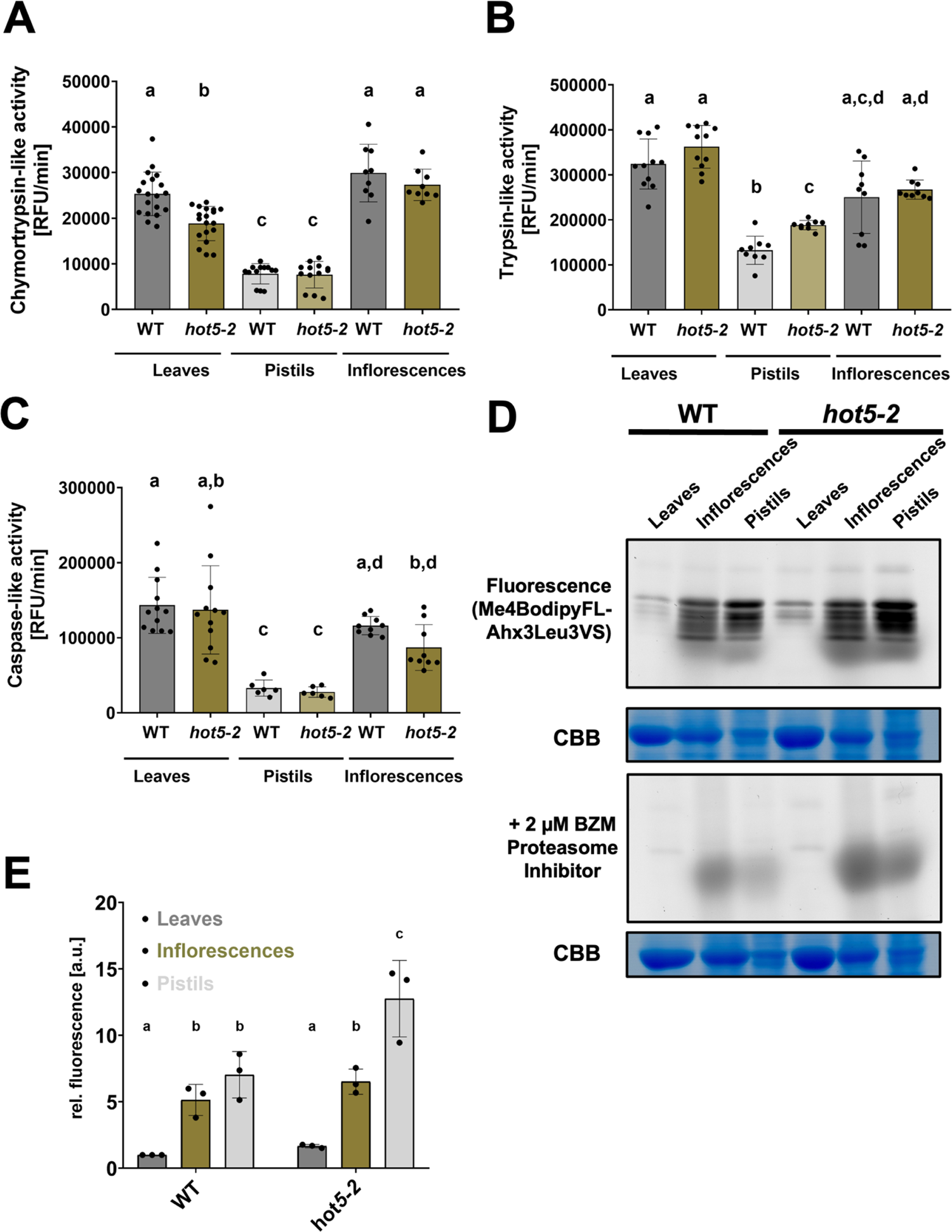
Proteasomal activity of different *A. thaliana* tissues. Chymotrypsin-like (A), trypsin-like (B) and caspase-like (C) activity of WT and *hot5-2* leaves, pistils and inflorescences were measured in the presence or absence of the proteasome inhibitor MG132. The fluorescence signal emitted by AMC cleavage was measured in a microplate reader via Gen5 software, averaged using Microsoft excel, and plotted using Graphpad Prism. Proteasome activity is shown after subtracting the RFU/min value observed in the reactions with MG132 and without protein. All experiments were repeated with three biological replicates, and the plotted data show the mean of at least two technical replicates. Error bars represent standard deviation and different letters indicate groups of significant differences at *p* ≤ 0.05 calculated by ANOVA with post hoc Tukey HSD. (D) In-gel fluorescence scans showing representative proteasome activity profiles using Me4BodipyFL-Ahx3Leu3VS as a substrate without (upper) or with (lower) the proteasome inhibitor bortezomib (BZM). The assay was repeated three times with three biological replicates for each tissue and genotype. (E) Relative quantification of fluorescence signals in comparison to the WT leaves samples obtained from (D) using ImageJ. Plotted are mean values with standard deviation. Different letters indicate groups with significant differences at *p* ≤ 0.05 calculated by two-way ANOVA with post hoc Tukey HSD.

## 4 Discussion

PTMs play crucial roles in regulating protein function and signaling pathways within biological systems. Among these modifications, S-nitrosation, the covalent attachment of a NO-group to cysteine residues of proteins, has emerged as a significant regulatory mechanism in many cellular processes in plants, including growth, development and photosynthesis as well as the general stress response (Wani et al., 2021; Gupta et al., 2022; Aranda-Caño et al., 2024). Understanding the dynamics of S-nitrosation and its impact on protein function requires reliable methods for detecting this PTM. The Biotin-Switch assay (BSA) has been the major method used for detection of S-nitrosated proteins (Forrester et al., 2009). The BSA involves several key steps, including blocking of free thiols to prevent non-specific S-NO detection, followed by the reduction of R-SNOs by ascorbate to expose the cysteine residues, and biotinylation of the exposed cysteines for isolation of the modified proteins. However, reports question the specificity of the ascorbate-dependent reduction of S-nitrosoproteins. It could be shown that ascorbate reduces protein disulfides of tubulin, tau and microtubule associated protein-2 that were isolated from porcine brain samples (Landino et al., 2006). In addition, the sulfenic acid form of yeast 1-Cys peroxiredoxin can be reduced by ascorbate (Monteiro et al., 2007). Therefore, using higher concentrations of ascorbate could potentially increase false-positives outcomes, whereas insufficient ascorbate concentrations would limit recovery of nitrosated proteins.

We used a direct enrichment technique, the MRC method, that uses organomercury to capture S-nitrosated proteins from complex biological samples. R-SNO proteins react with the organomercury compound on the resin, forming a stable mercury-thiol bond. A key advantage of the MRC method is its specificity towards S-nitrosated proteins, which bypasses the problematic ascorbate reduction step. Release of peptides from the resin also generates sulfonic acid, allowing direct identification of the nitrosated Cys residue. As with other methods, the MRC can be coupled with downstream analyses, such as Western blotting and mass spectrometry, allowing for the identification of modified proteins.

Previous R-SNO studies in plants included analysis of *A. thaliana hot5-2* mutant with higher endogenous NO levels (Hu et al., 2015) or the external application of NO donors (Lindermayr et al., 2005; Romero-Puertas et al., 2007; Hu et al., 2015). In contrast to the hundreds of S-nitrosated proteins/peptides typically identified by BSA-based approaches, the MRC method detected 919 S-nitrosated proteins and 533 R-SNO peptides (1102 protein groups in total) in a single experiment with triplicate biological replicates analyzed by mass spectrometry. The absence of previously identified S-nitrosated proteins in our analysis may be due to their extremely low abundance, differences in nitrosation resulting from endogenous NO versus exposure to exogenous NO donors, or regulatory differences in S-nitrosation in inflorescences as compared to seedlings and cell cultures. Results from our experiments are comparable to the MRC applied to mammalian systems. An MRC study of various mouse tissues identified 1011 S-nitrosocysteine residues in 647 proteins (Doulias et al., 2013), and the same group found 192 endogenously modified proteins with 328 R-SNO peptides in WT mouse liver (Doulias et al., 2010). In addition, another group used the MRC to analyze the specific R-SNO formation of PRXD2 in mammalian brains (Svistunova et al., 2019). In total, our data demonstrate the high sensitivity and reliability of the MRC method for conducting S-nitrosoproteomic analysis in plants.

Combining our data with previous work, more than 1650 S-nitrosated proteins have been identified in *A. thaliana*, providing one of the most comprehensive S-nitrosoproteome data sets for any organism. The MRC method together with quantitative label-free proteome profiling of specific reproductive tissues allowed us to identify the in-depth S-nitrosoproteome of floral tissues, which have not previously been analyzed. The identification of over 1000 S-nitrosated proteins in *hot5-2* mutant floral tissues is consistent with the elevated concentration of RNS in this mutant (Lee et al., 2008). We identified known S-nitrosation targets, including PrxIIE, GAPDH and multiple photosynthesis-related proteins, further highlighting the importance of NO homeostasis in plant processes such as photosynthesis as well as the general stress response. The discovery of hundreds of additional floral-specific S-nitrosated proteins offers avenues for future investigations of the biological role of S-nitrosation in plant growth and reproduction.

It is essential to note some limitations and considerations associated with the MRC technique. Similar to the BSA, proper controls and complete blocking of reduced thiols prior to R-SNO enrichment are important to prevent false-positive results. Additionally, the use of mercury-containing compounds raises environmental and safety concerns, necessitating proper handling and disposal procedures. In addition, *in hot5-2* samples we only detected site-specific R-SNO peptides (elution 2) for approximately 24% of the 919 S-nitrosated proteins recovered as bound to the resin (elution 1). It is unclear which step in the procedure results in this difference in peptide recovery, and further optimization is warranted. Previous MRC studies with mammalian cells only reported results of peptides from elution 2, providing no basis for comparison. A major drawback of any S-nitrosoproteome study is the masking of low-abundant S-nitrosated proteins by highly abundant proteins. Ribulose-1,5-bisphosphate carboxylase/oxygenase (RUBISCO) is one of the most abundant proteins in plants and has been reported to be S-nitrosated (Abat and Deswal, 2009). We applied a targeted mass exclusion for peptides associated with RUBISCO during LC-MS/MS analysis, but combining this approach with RUBISCO-depletion strategies might further improve the identification of more low-abundant S-nitrosated proteins and peptides.

Plants lacking GSNOR exhibit a severe fertility defect correlated with hormonal imbalance and increased callose accumulation in ovules, with ultimately reduced seed yield (Wang et al., 2024). Based on our quantitative proteomics and S-nitrosoproteomics data, we identified that many UGT proteins are S-nitrosated and differentially regulated in various *hot5-2* tissues. UGT proteins play crucial roles in plant hormone regulation that catalyze the transfer of sugar moieties to various substrates, including phytohormones like auxins, cytokinins and gibberellins (Ross et al., 2001). The importance of UGT S-nitrosation or of how nitro-oxidative modifications may impact UGT function remains unknown. However, the S-nitrosation and upregulation of UGTs in *hot5-2* plants makes them interesting targets for further investigation of their modulation by RNS and their importance in fertility.

In contrast, proteins belonging to the AGO protein family are less abundant in *hot5-2* pistils. AGO5 and AGO9 have been reported to play crucial roles in early female reproductive development (Olmedo-Monfil et al., 2010; Tucker et al., 2012), and S-nitrosation of AGO4 at Cys 482 is implicated in DNA methylation during stem cell regulation in *A. thaliana* (Zeng et al., 2023). The same study indicates that AGO4 can be S-nitrosated at multiple Cys residues besides Cys482, and we identified Cys342, 404, 457, 580 and 871 as additional S-nitrosation sites (Supplemental data 1, Tab 3). Protein sequence alignment shows that Cys404 of AGO4, located in the PAZ domain, is the only S-nitrosated Cys residue that is conserved among all *A. thaliana* AGO proteins. Therefore, it might be that AGO 5 (Cys467) and 9 (Cys376) are targets for S-nitrosation, leading to selective degradation, which could explain the down-regulation observed in *hot5-2* pistils. Similar regulation has been reported for proteins such as GSNOR, where S-nitrosation of Cys10 results in GSNOR degradation via autophagy (Chen et al., 2020). It will be important to elucidate the impact of nitro-oxidative modifications on the activity and overall involvement of posttranscriptional regulation in the context of plant reproduction and fertility.

Our results also implicate protein S-nitrosation in regulating protein quality control in floral tissues, specifically in pistils. We identified both regulatory and core proteins of the 26S proteasome as targets for S-nitrosation in floral tissues and uncovered specific upregulation of UPS components. UPS trypsin-like activity was also increased in *hot5-2* pistils, although UPS caspase- and chymotrypsin-like activities were unchanged. In contrast, all UPS protease activities in leaves and SePels were the same in *hot5-2* and WT, indicating that S-nitrosation does not negatively impact UPS function in these tissues. Mutations of individual regulatory particle subunits in *A. thaliana* show a wide range of phenotypes. Mutation in the RPN10 gene has been linked to hypersensitivity to ABA (Smalle et al., 2003), and a RPN12A T-DNA insertion allele exhibits reduced root elongation, delayed skotomorphogensis and altered growth responses to exogenously applied cytokinins (Smalle et al., 2002). The inactivation of RPN1A results in embryo lethality (Brukhin et al., 2005), and mutation in RPN5A impairs embryo and seed development (Book et al., 2009). The same authors conclude that this phenotype is likely due to a decreased stability of the holoproteasome complex. We found that RPN5A is S-nitrosated at Cys397 in floral tissues, and significantly upregulated in *hot5-2* pistils. Cys397 is located in the PCI domain that mediates and stabilizes protein-protein interactions, suggesting that NO might affect overall complex integrity or is involved in differential degradation of ubiquitinated target proteins. However, since the fertility defect observed in *hot5-2* is affecting early female gametophyte development, the precise role of RPN5 needs to be further evaluated.

There are only a few reports on the importance of NO on protein quality control. In contrast to our observations, rat vascular smooth muscle cells treated with various NO donors such as S-nitroso-N-acetylpenicillamine (SNAP) or GSNO showed a significant inhibition of all three (chymotrypsin-like, trypsin-like, caspase-like) UPS activities (Kapadia et al., 2009), suggesting that nitro-oxidative modifications impact the 26S core proteasome differently in different organisms and tissues. Besides the direct effect on the 26S proteasome, NO has been suggested to also interfere with the ubiquitin conjugation system. For instance, S-nitrosation of the RING-finger E3 ligase parkin inhibited its activity (Tsang et al., 2009). Moreover, NO can trigger the ubiquitin-mediated proteasomal degradation of S-nitrosated proteins in plants, as reported for ascorbate peroxidase 1 (APX1) and ABA insensitive 5 (ABI5) (De Pinto et al., 2013; Albertos et al., 2015). All these results highlight the potential role of NO in regulating the UPS. However, although our data point to a molecular link between NO signaling in floral tissues and protein quality control, many open questions remain. It will be of interest to understand if the protein composition of the proteasome is different in reproductive tissues, if there are specific UPS targets for degradation in floral tissues, and the precise role of S-nitrosation in UPS regulation.

In conclusion, the organomercury-resin capture of S-nitrosated proteins is a valuable method for the specific isolation and analysis of proteins carrying this biologically significant protein modification. Its application contributes to advancing our understanding of the roles played by S-nitrosation in various cellular pathways and developmental processes. In particular, we found S-nitrosation of multiple UPS components in reproductive tissues, revealing a potential mechanism linking protein quality control to fertility defects and deregulated NO homeostasis. Further understanding of the relationship between GSNOR, NO homeostasis, and plant fertility may lead to new approaches to crop improvement through increased fertility.

## 6 Author Contributions

Patrick Treffon designed and performed research, analyzed data, and wrote the paper. Elizabeth Vierling: designed and supervised the project, analyzed data, and wrote the paper. Both authors approved the final manuscript.

## Acknowledgments

We thank Drs. Harry Ischiropoulos, Jordanis Zakopoulos and Paschalis-Thomas Doulias for taking their time and care to provide us with the opportunity to learn the MRC technique from them. We also would like to thank Dr. Steve Eyles for his help with mass spectrometry and Dr. Eric Strieter for providing us with the proteasomal activity probe and bortezomib. Mass spectral data were obtained at the University of Massachusetts Mass Spectrometry Core Facility, RRID:SCR_019063.

## 8 Supplementary Material

Supplemental data set 1. List of S-nitrosated proteins

Supplemental data set 2. MapMan classification of S-nitrosated proteins

Supplemental data set 3. KEGG enrichment analysis

Supplemental data set 4. List of differentially regulated proteins in pistils

Supplemental data set 5. List of differentially regulated proteins in SePels

Supplemental data set 6. List of differentially regulated proteins in leaves

Supplemental data set 7. List of identified UGT proteins

Supplemental data set 8. Correlation analysis RSNO vs. quantitative

## 12 Data Availability Statement

The mass spectrometry data are accessible in the MassIVE repository (https://massive.ucsd.edu) with the dataset identifiers (MSV000094148, S-nitrosoproteome all proteins; MSV000095464, S-nitrosoproteome site-specific; MSV0000094145, pistil proteome; MSV000094146, SePel proteome) and password hot5-2.

---

**Figure S1.** Control treatments demonstrate the specificity of the MRC method for R-SNO identification. Increasing total protein amounts of *hot5-2* leaves (5 to 20 mg) treated with or without 1 mM ascorbate (ASC)/100 µM CuSO_4_ or UV light (photolysis) were subjected to the MRC method, and SDS-PAGE-separated proteins were visualized by silver-staining. Control treatments effectively eliminate MRC capture.

**Figure S2.** Functional characterization of S-nitrosated proteins. MapMan classification of the 1102 S-nitrosated proteins identified as enriched in *hot5-2* and WT inflorescences based on metabolic-(A) and regulatory pathways (B). Each red square represents a S-nitrosated protein for a given pathway. Proteins from the MapMan output are listed in the Supplemental Data Set 2.

**Figure S3.** Volcano plots of differentially expressed AKR4C proteins. The pistil and SePel dataset are reported in this study, whereas the leaf dataset was published previously (Treffon et al., 2021). AKR4C Proteins (PFAM ID PF00248) in different *hot5-2* and WT tissues are shown. UniProt identifier: AKR4C8, O80944; AKR4C9, Q0PGJ6; AKR4C10, Q84TF0; AKR4C11, Q9M338. The -log10 of the corrected *p*-values is plotted against the log2-fold change (FC) in protein levels. FDR cutoff (5%; S0= 0.1) is indicated by black lines. A list of the significantly regulated proteins for each tissue is provided in the Supplemental Data Sets 4-6.

**Figure S4.** Organ samples used for quantitative proteome analysis. Pistils (Ps), sepals (S) and petals (P) of WT and *hot5-2* flowers at flower development (FD) stages 12-13 were used.

**Figure S5.** Summary of 26S proteasomal proteins that are S-nitrosated in inflorescences (black) and up-regulated in pistils of *hot5-2*. A list of all upregulated and S-nitrosated UPS proteins can be found in Table 2 and Supplemental Data Set 1 and 4. Eukaryotic 26S proteasomes consist of a 20S core and one or two 19S regulatory caps. The 19S regulatory particles are involved in the recognition and unfolding of ubiquitinated substrates, while the catalytic activity takes place in the 20S proteasome. The 20S proteasome is a barrel-shaped protein complex composed of four stacked rings that consist of seven subunits each. The two outer base-rings are made up of α-subunits and interact with the 19S regulatory caps. The two inner rings consist of β-subunits that are responsible for proteolysis.

